# Biological aging and comorbidities are linked to HIV reservoir size despite suppressive antiretroviral therapy

**DOI:** 10.64898/2026.11.09.765455

**Authors:** Mareva Delporte, Twan Otten, Kavita Mehta, Evy E. Blomme, Sofie Rutsaert, Maxime Verschoore, Victoria Rios Vazquez, Nadira Vadaq, Vasiliki Matzaraki, Casper Rokx, Janneke Stalenhoef, Marvin Berrevoets, Jan Van Lunzen, Yubo Zhang, Leo Joosten, Mihai G. Netea, Jéssica C. dos Santos, Sarah Gerlo, Wim Trypsteen, André van der Ven, Linos Vandekerckhove

**Affiliations:** HIV Cure Research Center, Ghent University, Ghent, Belgium; Radboud University Medical Center, Nijmegen, The Netherlands; ETZ Tilburg, The Netherlands; Erasmus University Medical Center Rotterdam, The Netherlands; OLVG, Amsterdam, The Netherlands; Department of Immunology and Metabolism, Life and Medical Sciences Institute, University of Bonn, Bonn, Germany; The Ghent University Digital PCR Center, Ghent, Belgium

## Abstract

**Background:** The HIV reservoir persists in long-living immune cells after antiretroviral therapy initiation. Proviral DNA integrates at preferential host DNA sites, while HIV transcription continues under treatment, with potential biological consequences. Treated people with HIV remain at increased risk for premature aging and non-AIDS comorbidities compared to people without HIV. Whether HIV reservoir size and activity impact health outcomes during treatment remains unclear.

**Methods:** We analyzed 1895 people with HIV using antiretroviral therapy from the 2000HIV cohort to study associations between the reservoir and historic and prospective virologic and immunological response markers, comorbidity outcomes and biological aging scores. The reservoir was assessed by quantifying total and intact proviral HIV DNA in circulating CD4+ T cells, along with measures of HIV transcriptional activity, including cell-associated HIV RNA.

**Results:** HIV reservoir markers were highly correlating and associated with older aging scores and adverse virologic outcomes, including quantifiable and residual viremia at study entry and during follow-up. Total HIV DNA, unlike the minor fraction of intact HIV DNA, correlated with prevalent cardiovascular disease, subclinical atherosclerotic disease, liver fibrosis and incident malignancies independent of circulating inflammatory markers and lowest CD4 nadir. Total HIV RNA associated with prevalent cardiovascular disease and liver fibrosis.

**Conclusion:** The HIV reservoir associated with older aging scores and long-term health outcomes in people with HIV using antiretroviral therapy, independent of inflammation and immune suppression history. Whether reservoir-reducing interventions lower the risk of non-AIDS comorbidities should be studied in people with high reservoir markers.

**Funding and trial registration:** ViiV Healthcare; 2000HIV ClinicalTrials.gov number: NTC03994835

**Graphic summary:** 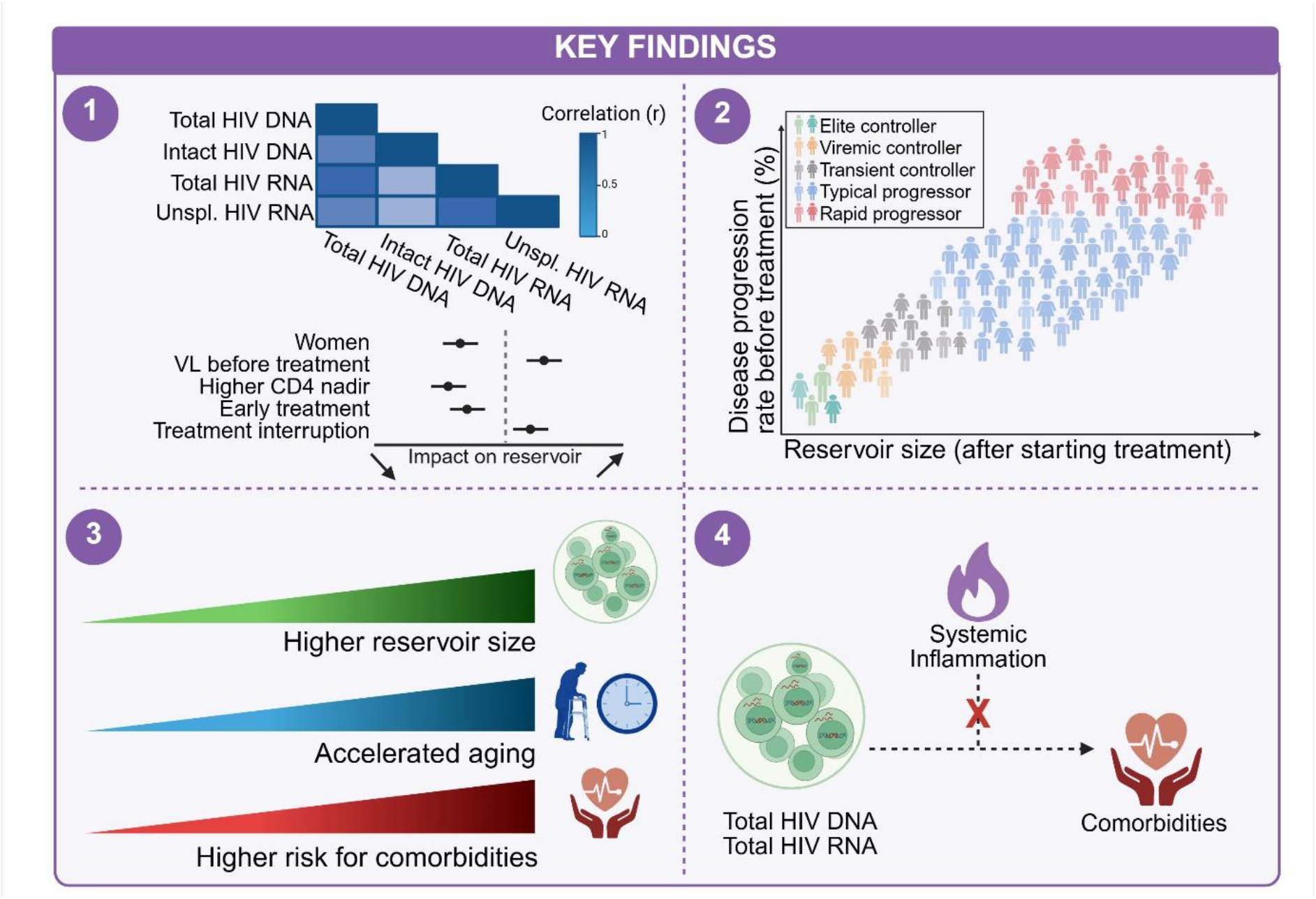

**Highlights:**

- HIV reservoir markers in circulating CD4 cells, including total and intact HIV DNA as well as total and unspliced HIV RNA, are highly correlated and show largely overlapping clinical and immunological characteristics.
- People with less favourable HIV clinical phenotypes before ART show higher levels of all measured HIV reservoir markers.
- Higher total HIV DNA and total HIV RNA associate with older biological aging scores and higher non-AIDS comorbidities risk after start ART.
- The effect of the reservoir on comorbidities is not purely dependent on systemic inflammation

## Introduction

Type 1 human immunodeficiency virus (HIV) infects and kills CD4 T cells, which ultimately leads to severe immunodeficiency and death after a median period of 10-13 years, unless antiretroviral therapy is given [1, 2]. Not all people with HIV progress in such a way: HIV controllers can spontaneously suppress HIV replication and can therefore maintain high CD4 cell counts with a favorable prognosis without antiretroviral therapy [3]. In contrast, rapid progressors with high plasma viral loads and fast CD4 cell decline will develop severe immunodeficiency within 3-5 years after infection [2]. Since the introduction of combination antiretroviral therapy, HIV infection has shifted from a fatal diagnosis to a chronic condition needing permanent treatment [4]. Over time, it became evident that early initiation of antiretroviral therapy significantly improves long-term outcomes with life expectancies that reach or equal the HIV uninfected population [5, 6]. In contrast, delayed treatment and low CD4 cell counts, concerning 30-70% of people with HIV in certain populations [7, 8], increase the risk for comorbidities such as cardiovascular diseases and cancers, with persistent immune activation and inflammation as reported important drivers [5, 6, 9]. Such immune imbalance is especially evident in immunological non-responders, who fail to completely restore CD4 cell counts despite suppressive antiretroviral therapy, and show a high risk for premature death and comorbidities [10].

Apart from immune dysregulation, HIV itself, especially the HIV-1 reservoir, may increase comorbidity risk. This reservoir primarily consists of a pool of long-lived, resting CD4 T cells harbouring HIV genomes [11–15] that consists of mostly defective proviruses and a minor fraction of intact, replication-competent proviruses [16]. Proviral integration in host DNA is not random, but occurs at preferential sites, which may have biological consequences for the host but may also promote clonal expansion of the reservoir [17]. Furthermore, the host may also be affected by products of HIV transcription that persists under antiretroviral therapy. Clinical effects of the reservoir were reported in a small-scale study that showed an association between a larger HIV reservoir size and plaque presence, independent of inflammation [18], however, large-scale studies are lacking.

Reservoir assessment remains challenging because of complex and expensive technologies. Therefore only a few studies have included a large number of participants [19, 20]. Data from the Swiss and RID-HIV cohorts showed in 1057 and 892 participants, respectively that early treatment, low viral zenith and high CD4 nadir associated with lower total HIV-1 DNA reservoir levels [19]. Reservoir data from extreme phenotypes come from smaller cohort studies: spontaneous HIV controllers harbour a relatively small reservoir [21, 22] in contrast to immunological non-responders with larger reservoir sizes [23].

The 2000HIV cohort study included 1895 virally suppressed people with HIV from four Dutch HIV-treatment centers, with one of the aims to study the impact of the latent HIV reservoir on health outcomes [24]. We quantified total HIV DNA, intact HIV DNA, defect HIV DNA, total HIV RNA and unspliced HIV RNA in circulating CD4 T cells and performed high-throughput plasma proteomics. Furthermore, we documented the complete medical history, assessed comorbidities using imaging techniques at baseline, and collected clinical data during a 2-year follow-up. We first compared reservoir parameters across HIV phenotypes, then examined their association with both historic and prospective virological, immunological and clinical outcomes. Finally, the plasma proteome was used to calculate proteomic aging scores and to check whether associations with comorbidities were driven by systemic inflammation markers.

## Methods

### Sex as a biological variable

The study population comprised predominantly male individuals, reflecting the sex distribution among people living with HIV in the Netherlands. Sex assigned at birth was included as a covariate in statistical analyses where appropriate.

### Study population and procedures

This prospective cohort study is based on the 2000HIV study (NTC03994835) that included 1895 asymptomatic people with type 1 HIV who were virally suppressed because of antiretroviral therapy or spontaneous control. Recruitment took place between October 2019 and October 2021 at HIV treatment centers in the Netherlands [24]. At enrollment, complete medical history and medication use were recorded using structured questionnaires and electronic patient records, supplemented and cross-checked with data from the ATHENA cohort (Stichting HIV Monitoring). Assessments included documentation of AIDS and non-AIDS comorbidities (e.g., NAFLD, cardiovascular disease), carotid ultrasound and FibroScan® (in a subgroup). After medical examination during the first visit, blood was sampled for multi-omics and reservoir assessment. At the two-year follow-up visit, incident events were reassessed using the same structured questionnaire and electronic patient records. Antiretroviral therapy adherence was assessed using MASS-8 questionnaire, and extracting suspected non-adherence when documented by the treating physician in electronic patient record. The supplement methods include operational definitions and HIV phenotype definitions (Table S1).

Approval for the study was granted by the Independent Review Board Nijmegen (ref. NL68056.091.81). All participants provided written informed consent. The study was funded by VIIV Healthcare, which was not involved in any analysis nor writing of the manuscript.

### Quantification of HIV DNA and HIV RNA from CD4 T cells

Total and intact HIV-1 DNA levels were assessed in CD4 T cells by digital PCR (dPCR) using the Rainbow proviral HIV-1 DNA assay [10]. CD4 T cells were enriched from 40 million cryopreserved PBMCs by negative selection using the EasySep Human CD4 T cell isolation kit on the Robosep-S (Stemcell Technologies, Vancouver, Canada). Cells were automatically counted using the Countess 3 Automated Cell Counter (Thermofisher Scientific, Massachusetts, United States) before dry pelleting. Genomic DNA (gDNA) was extracted using the QiaAmp DNA mini kit on the Qiacube (Qiagen, Hilden, Germany) using two elution steps of 50 µL. DNA concentrations were measured using the SpectraMax i3x using 2 µL eluted DNA with the SpectraMax Quant AccuBlue HiRange dsDNA assay Kit (Molecular Devices, San Jose, California, United States) after which samples were stored at −20 °C. Quantification was performed in triplicate using the Rainbow proviral HIV-1 DNA assay on the QIAcuity Four platform (Qiagen, Hilden, Germany). For samples with DNA concentrations below 90 ng/µL, 18 µL of DNA was used per replicate. For samples with DNA concentrations above 90 ng/µL, 10 µL was used. HIV-1 DNA levels were normalized by measuring the reference gene RPP30 in duplicate. For normalization and DNA shearing assessment, a 100-fold dilution was made for each sample from which 5 µl was used as input.

The RU5 region from the Rainbow assay was used to assess total HIV-1 DNA levels. Intact proviruses were obtained by assessing the presence of at least two (psi and env) and maximum 5 target regions (RU5, psi, gag, pol and env) in the Rainbow assay. The Rainbow Shiny tool was used for automatic thresholding, and this was manually adapted per sample when a threshold crossed the double-positive population. For individuals for which the intactness value was 0, the results for the individual targets were manually reviewed. If positive partitions were observed in all target regions, the result was judged as undetectable, and the intact reservoir was manually set to one copy in the total cell input for the sample. If no positive partitions were detected for the psi and/or env region, no intactness level was reported as this was deemed to be due to a signal failure.

When sufficient cells were counted, cells were pelleted and RNA was extracted using the RNeasy plus mini kit on the Qiacube (Qiagen, Hilden, Germany), using 30 µL to elute. RNA concentrations were measured using the broad range RNA concentration kit on the Qubit, ranging between 4 and 168 ng/µL. RNA was immediately reverse transcribed into cDNA in a 31.75 µL reaction mix using the Superscript III kit [25 µL sample, 1.6 µL of 10x SuperScript III buffer, 1.6 µL of 50 mM MgCl2, 0.8 µL of random hexamers (50 ng/μL; Invitrogen), 0.8 μL of 50 μM poly-dT15, 0.8 μL of 10 mM deoxynucleoside triphosphates (dNTPs), 0.4 μL of RNAseOUT (40 U/μL; Invitrogen), and 0.8 μL of SuperScript III RT (200 U/μL; Invitrogen)]. The reverse transcription program consisted of 10 min at 25 °C, followed by 50 min at 50 °C and 5 min at 85 °C. Samples were stored at −20 °C until further measurements. HIV-1 RNA levels were measured in duplicate by dPCR using the triplex HIV-1 RNA dPCR assay on the QIAcuity Four platform using 12 µL per replicate (Qiagen, Hilden, Germany). HIV-1 RNA was normalized based on cell number in the pellet and qPCR of housekeeping genes GAPDH and HPRT1, for which 2.5 ng cDNA was used as input and were run in duplicate. Samples that did not meet the quality control criteria (abnormal cDNA volume after storage, dPCR partitioning rate < 75% or qPCR ΔCq between replicates higher than 1.5) were excluded for further analysis.

### Proteomic profiling

Proximity extension technology (Olink® Explore 3072 panel) was used to assess the plasma proteomic profile as described earlier [24, 25]. Results were reported as Normalized Protein Expression (NPX) values after quality control and normalization. In addition to the Olink PEA QC process, proteins were excluded when the target protein was detected in less than 75% of the samples. In total, 2367 proteins were included for follow-up analysis.

### Biological age calculation

Biological age scores were calculated by applying bootstrap LASSO models to the expression levels of specific proteins depending on the aging score, as described in Zhang et al. [26].

### Statistical analysis

Unreportable values from reservoir quantification were excluded from analysis as they could be due to technical issues, as well as values from participants on immunosuppressants, leaving 1822 participants with total HIV DNA levels, 1402 participants with intact/defect HIV DNA levels, 1428 with total HIV RNA levels and 1398 with unspliced HIV RNA levels. First, we summarize the reservoir size across the complete cohort and predetermined HIV phenotypes. Those participants who did not satisfy the criteria for a phenotype were excluded from the respective analyses (Table S1).

Second, univariable and mutlivariable linear regression analyses were conducted to identify factors associated with the different reservoir markers (total HIV DNA, intact HIV DNA, total HIV RNA and unspliced HIV RNA) in participants receiving antiretroviral therapy (n = 1798, n = 1379, n = 1411 and n = 1382 respectively). The variables included clinical and immunological parameters that were not assessed as outcome variables.

Thereafter, associations between reservoir size and clinical outcomes were tested in participants receiving treatment using (multiple) logistic regression, reporting odds ratios (OR) with 95% confidence intervals (CI) per 1 log_2_ increment in reservoir size. Variables with p < 0.05 in univariable analysis were entered into mutlivariable models adjusting for age, sex, and CD4 nadir. Sensitivity analyses tested the independence of observed associations from other preselected variables relevant to the comorbid condition, known or suspected inflammatory mediators implicated in HIV-related inflammation (sCD14, sCD163, CXCL10, IFNγ, IL10, TNFα, IL1β, IL6).

Additionally, the Youden index was used to determine optimal total HIV DNA cut-off values, stratifying participants into high- and low-risk groups for each outcome, maximizing the balance of sensitivity and specificity.

Associations between HIV reservoir markers and biological aging scores were evaluated using multivariable linear regression, adjusted for chronological age, sex at birth, and ethnicity. Multiple testing was controlled using the Benjamini-Hochberg false discovery rate procedure. Regression coefficients with 95% confidence intervals are reported.

## Results

### Cohort and reservoir characterization

The study population in which the reservoir was assessed (n = 1822) was mostly representative of a Western European HIV population with a median (IQR) age of 53 (44-60) years; 270 (15%) women; 443 (24%) non-white participants and 1315 (71.8%) men who have sex with men (MSM) (Table S2, Figure S1). Extreme phenotypes included 104 spontaneous controllers, 80 rapid progressors, and 87 immunological non-responders. Apart from 24 spontaneous controllers, antiretroviral medication was initiated in all participants.

Total proviral HIV DNA levels were successfully obtained in 1822 participants, with a median of 603 cp/Mio cells [IQR: 251, 1247] (Figure 1a). Intact and defective proviral HIV DNA levels were obtained in 1402 participants, with a median of 16 intact cp/Mio cells [IQR: 4, 51], of which 450 with undetectable intactness levels (Figure 1b, Figure S2). Defective proviral DNA levels resulted in a median of 547 defect cp/Mio cells [IQR: 208,1160] (Figure 1c). Total HIV RNA levels and unspliced HIV RNA levels were obtained in 1428 participants and 1399 PLWH with a median of 122 [IQR: 33,374] (Figure 1d) and 17 cp/Mio cells [IQR: 3,62] (Figure 1e). All reservoir markers were highly correlating and interestingly, among the proviral DNA markers, defective HIV DNA correlated best with total HIV RNA (Pearson rho = 0.71, p < 0.001; Figure 1f).

**Figure 1:**
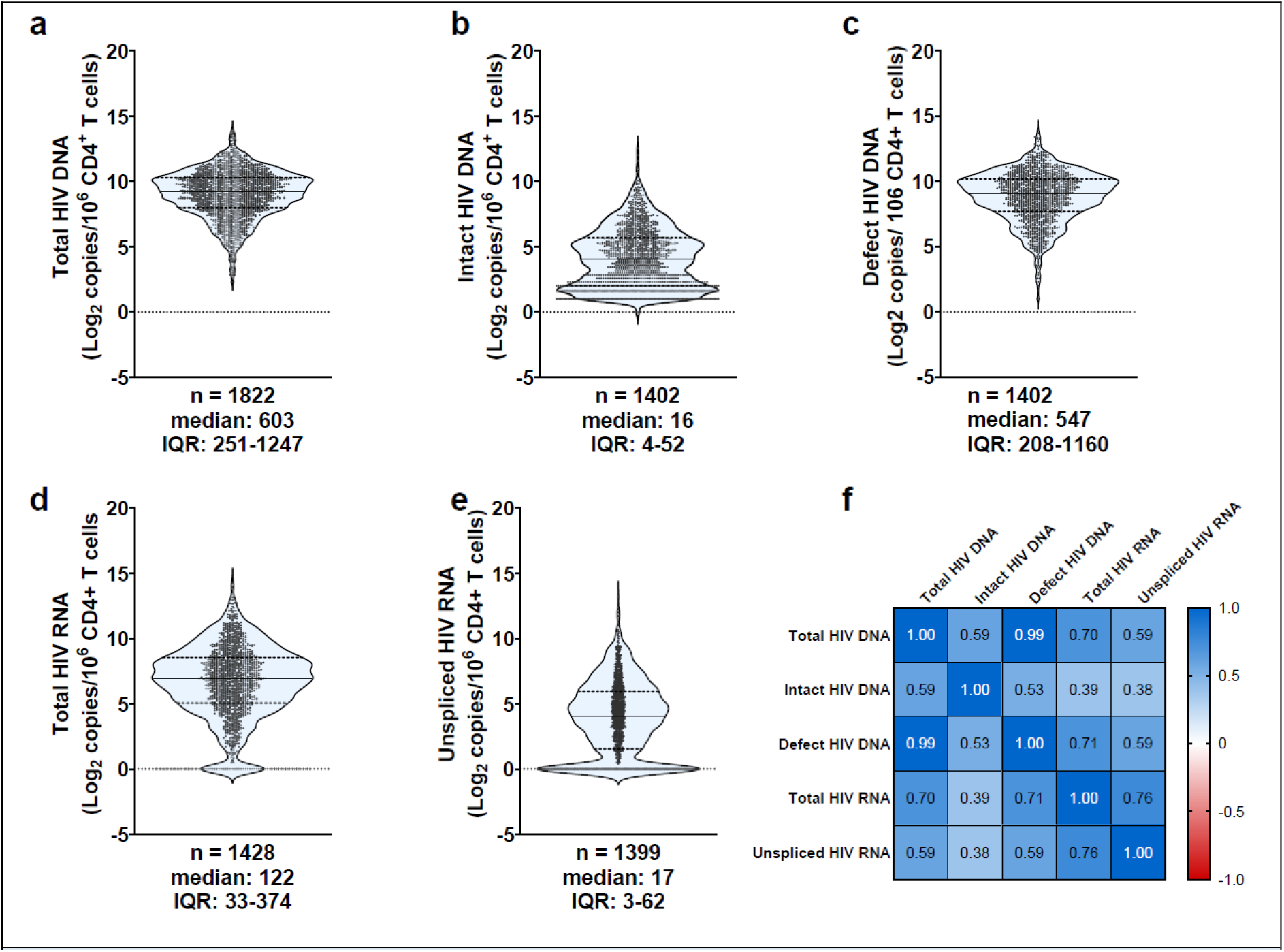
Reservoir composition in 2000HIV cohort. Violin plots showing the distribution of (a) total HIV DNA, (b) intact HIV DNA, (c) defective HIV DNA, (d) total HIV RNA, and (e) unspliced HIV RNA levels, expressed as log2 copies per 10^6^ CD4+ T cells. Horizontal dashed lines indicate the median and interquartile range (IQR). For each measure, the sample size (n), median, and IQR are reported below the corresponding plot. (f) Pearson correlation matrix showing correlations between HIV reservoir and transcriptional measures. Color intensity reflects the strength and direction of the correlation, and values within squares represent Pearson’s correlation coefficients.

### Reservoir across HIV phenotypes

People with HIV are heterogenous in terms of demographics, immune status, disease progression, treatment regimens and HIV duration making it likely that the HIV reservoir is also highly diverse [27]. Our data indeed show that the reservoir composition was significantly different across various HIV phenotypes (Figure 2). When stratifying PLWH based on disease progression and viral load prior to treatment initiation, distinct patterns emerged. Treatment-naïve elite controllers exhibited the smallest reservoir size, followed by viremic controllers. Interestingly, no statistically significant differences for any of the reservoir markers were observed between transient controllers, typical progressors and rapid progressors.

**Figure 2:**
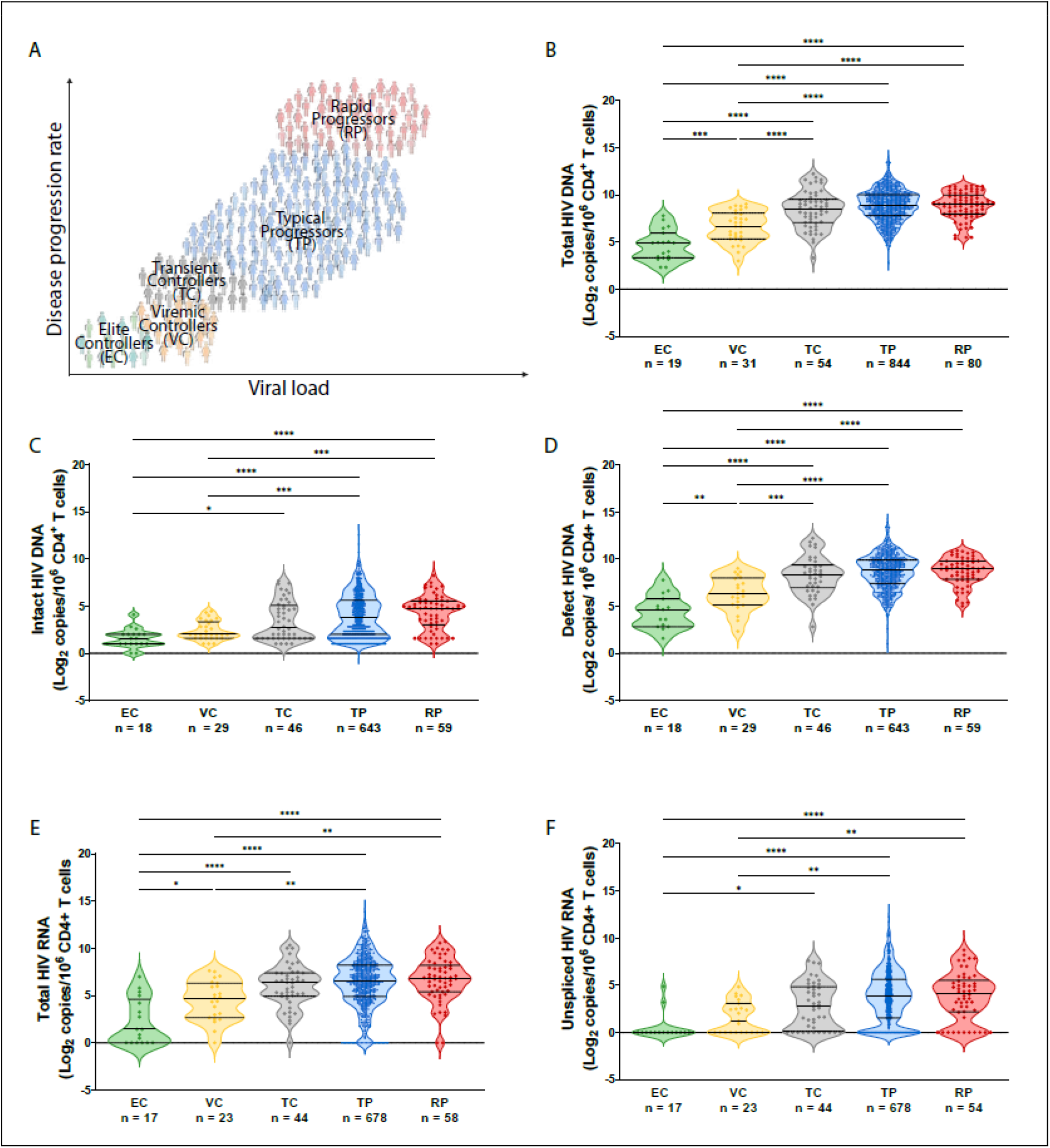
Reservoir composition across HIV phenotypes. (A) Schematic representation of HIV phenotypes according to viral load before starting treatment and disease progression rate, resulting in elite controllers (EC), viremic controllers (VC), transient controllers (TC), typical progressors (TP) and rapid progressors (RP). (B-D) Violin plots showing the distribution of (B) total HIV DNA, (C) intact HIV DNA, (D) defective HIV DNA, (E) Total HIV RNA and (F) unspliced HIV RNA. Horizontal dashed lines indicate the median and interquartile range (IQR). Sample sizes for each group are shown below the corresponding violins. Participants that did not meet criteria for certain groups, were not included in subgroup analysis. Stars indicate significance levels of Tukey’s multiple comparisons test.

### Sex and HIV related factors associate with HIV reservoir size

Results of univariable analysis exploring the relation between the reservoir markers and clinical data are presented in Figure 3 with a more extensive list of variables in supplement (Figure S3-6). Many of these variables that relate to the HIV reservoir, also strongly relate to each other (Figure S7), so should be carefully interpreted. Importantly, the observed differences in reservoir levels over time may reflect changes in HIV management guidelines rather than the direct impact of specific treatment regimens (Figure S8).

**Figure 3:**
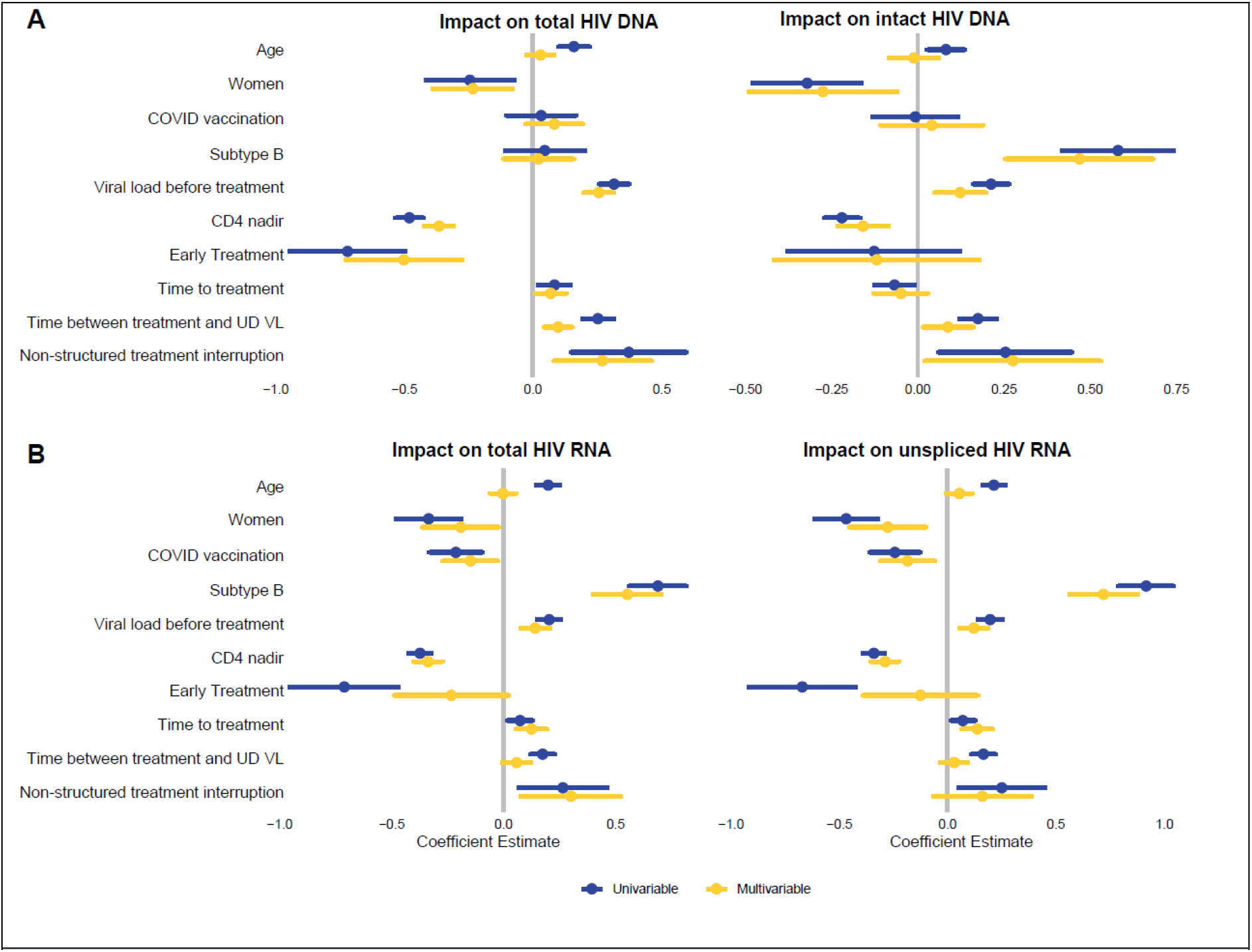
Determinants of HIV reservoir. Coefficient plots showing covariables associated with total HIV DNA levels, intact HIV DNA levels, total HIV RNA levels and unspliced HIV RNA levels in univariable and multivariable analysis. Dots represent coefficient estimates, bars represent 95% confidence intervals and stars indicate significance levels. (Subtype B: subtype B carriers versus non subtype B carriers; Viral load before Treatment: plasma HIV RNA copies/ul before start antiretroviral therapy; Non-structured Treatment interruption: non-structured antiretroviral therapy interruption for more than 3 months compared to no interruption; Time to Treatment = effect of time between HIV diagnosis and start antiretroviral therapy; Time Treatment to UD VL: effect of time between start antiretroviral therapy and undetectable plasma viral load)

Subsequent mutlivariable linear regression (Figure 3) using clinically relevant variables showed similar impact on the different reservoir markers. Women and those who initiated antiretroviral therapy during acute HIV infection exhibited lower reservoir levels, whereas higher reservoir levels are found in participants with non-structured treatment interruption, higher viral load before antiretroviral therapy, longer time between treatment initiation and undetectable plasma viral load or lower CD4 nadir. Participants with subtype B had higher intactness levels compared to non B subtypes as well as higher HIV RNA levels (Figure 3). Vaccination against COVID19 did not impact proviral HIV-1 DNA levels (Figure 3a) but was associated with lower total HIV RNA levels (Figure 3b).

### Associations of viral reservoir with virologic, immunologic, aging and comorbidity outcomes

To explore the associations of the reservoir with virologic, immunologic and clinical outcomes, we first conducted a univariable logistic regression analysis (Table S3-S7), after which we examined the independent effects of the reservoir on these significant outcomes adjusting for age, sex and CD4 nadir (Figure 4) and other known risk factors (Table S8-12).

**Figure 4:**
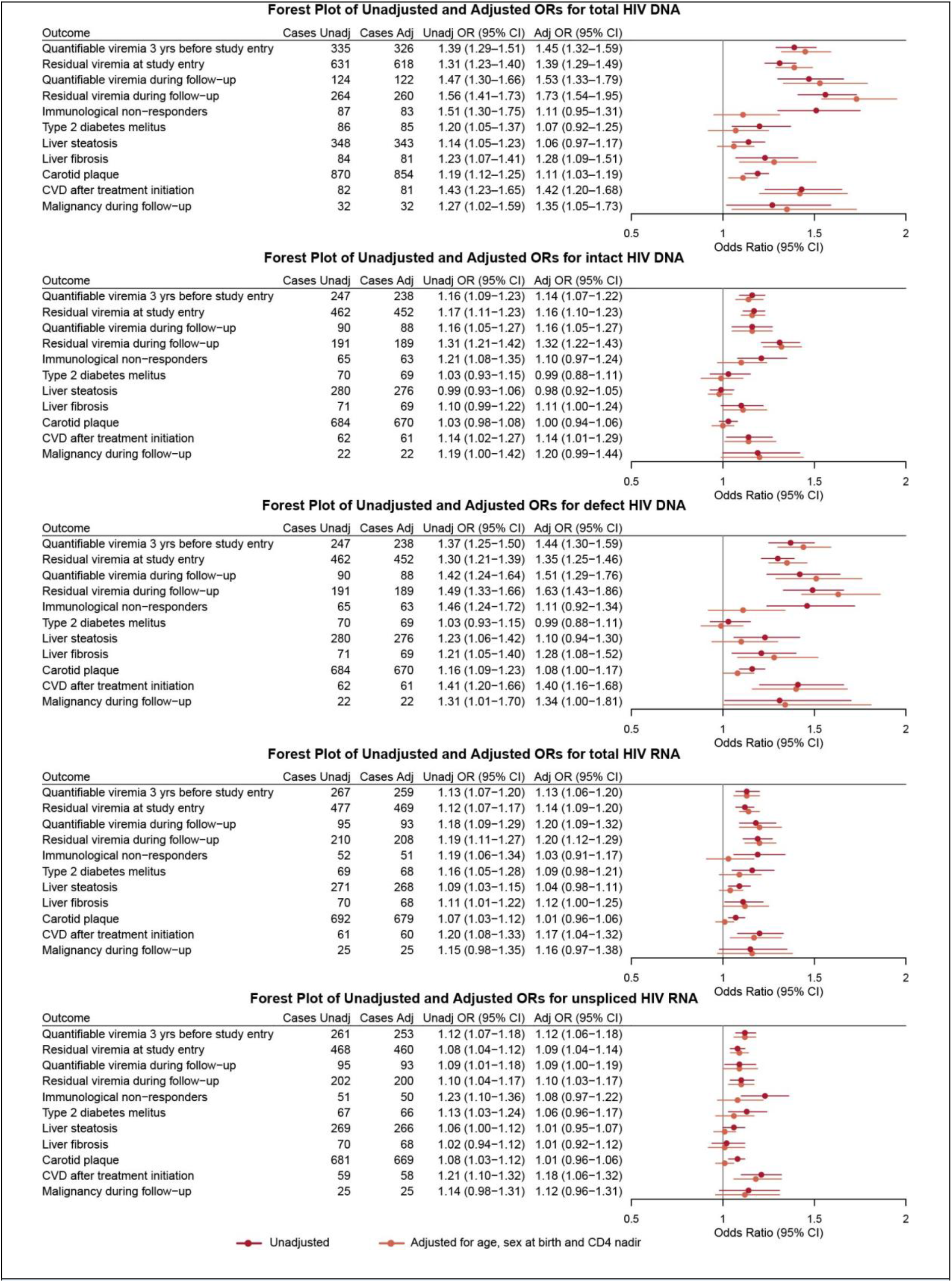
Forest plot of unadjusted and adjusted odds ratios related to virological or clinical outcomes. Univariate logistic model and multivariable logistic model with age, sex and CD4 nadir as covariates was used to determine the probability that reservoir markers relate to virological and clinical outcomes. Results are expressed as unadjusted odds ratios (OR) and adjusted odds ratios (aOR) per 1 log_2_ increment in reservoir size. Dots indicate the odds, vertical lines indicate confidence interval (CI95%) of the odds per clinical outcome. When 95%CI crosses the grey line, results are not significant. CVD= cardiovascular disease.

#### Viral reservoir size is associated with historic and prospective adverse plasma virologic endpoints

Larger total reservoir size was associated with higher odds of quantifiable viremia in the three years prior to cohort entry (aOR 1.45; 95% CI 1.32-1.59) and with residual viremia at entry (aOR 1.39; 95% CI 1.29-1.49; Figure 4a). Similar associations were observed for quantifiable viremia (aOR 1.53; 95% CI 1.33-1.79) and residual viremia (aOR 1.73; 95% CI 1.54-1.95) during the 2-year follow-up (Figure 4a). These findings remained robust after adjustment for treatment duration and time from HIV diagnosis to treatment initiation (Table S8), suggesting that larger viral reservoirs are linked to adverse plasma virologic outcomes, irrespective of disease progression before antiretroviral therapy, treatment delay, or treatment duration. Similar associations were observed for the intact/defect reservoir with both quantifiable viremia and residual viremia at cohort entry and during follow-up, as well as for HIV RNA markers (Figure 4b-e, Table S9-12).

#### Immunologic recovery is not associated with reservoir size

While all reservoir markers were associated with the odds of being an immunological non-responder (OR 1.19-1.51), this effect diminished after adjustment for age, sex and CD4 nadir (Figure 4). While the total viral reservoir moderately correlated with CD4 nadir (rho= −0.47; Figure S9), the size of the total reservoir itself was not independently associated with immunological recovery.

#### Viral reservoir size is associated with older biological aging scores

Next, we assessed whether viral reservoir size associated with biological aging scores, while accounting for chronological age, sex at birth and ethnicity (Figure 5) [26]. Interestingly, higher levels of cell-associated HIV RNA were associated with older biological aging scores across multiple domains, including conventional, immune, all-organ, organismal, artery and intestine age after correction for multiple testing. Higher total HIV DNA levels were associated with older conventional age and older all organ age. Together, these findings indicate that both HIV transcriptional activity and reservoir burden are associated with accelerated biological aging.

**Figure 5:**
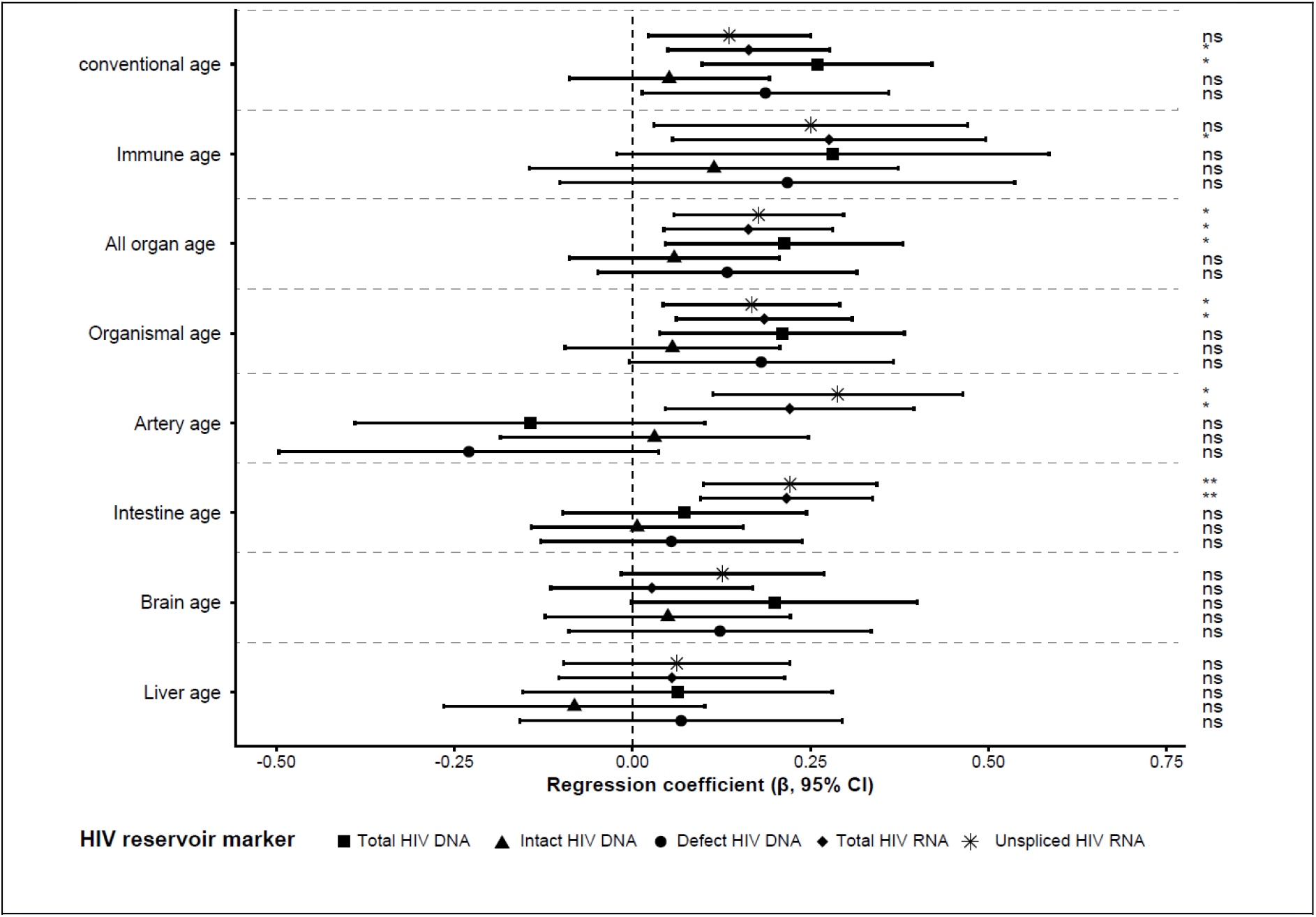
Forest plot showing regression coefficients (β) and 95% confidence intervals for the associations between biological age estimates (liver, brain, intestine, artery, organismal, all-organ, immune, and conventional age) and HIV reservoir markers, including defective proviruses (●), intact proviruses (▴), total HIV DNA (▪), total HIV RNA (◆), and unspliced HIV RNA (✳). Positive β values indicate higher biological age associated with higher HIV reservoir measures. Models were adjusted for age, sex, and ethnicity. The dashed vertical line indicates no association (β = 0). Statistical significance is denoted as not significant (ns), adj. P < 0.05 (*), adj. P <0.01 (**).

#### Both HIV DNA and HIV RNA associate with increased comorbidity

Total HIV reservoir significantly associated with different prevalent comorbidities (Figure 4a), such as liver fibrosis (aOR 1.28; 95%CI 1.09-1.51), presence carotid plaques (aOR 1.11; 95% CI 1.03-1.19) and cardiovascular disease (CVD) developed after treatment initiation (aOR 1.41; 95% CI 1.19-1.67; Figure 4a). The effects remained significant after additionally adjusting the cardiovascular outcomes for traditional risk factors (plaque: aOR 1.16; 95% CI 1.03-1.31, and CVD: aOR 1.64; 95% CI 1.31-2.04) and liver fibrosis for BMI, treatment duration, time between diagnosis and treatment initiation (aOR: 1.24; 95%CI 1.04-1.48; Table S7). Associations between total HIV DNA and liver steatosis and type 2 diabetes mellitus lost significance after adjusting for age, sex and CD4 nadir (Figure 4a).

During the 2-year study follow-up, larger total reservoir size was associated with new diagnoses of malignancy (aOR 1.35; 95% CI 1.05-1.73) and was not affected by further adjustment for years on antiretroviral therapy, time between diagnosis and treatment initiation, or prior malignancy history (Table S7). Interestingly, similar associations were observed between other reservoir markers, including defect HIV DNA and HIV RNA, and specific outcomes. These included cardiovascular disease developed after treatment initiation (defect HIV DNA aOR: 1.40; 95% CI 1.16-1.68, total HIV RNA aOR: 1.17; 95% CI 1.04-1.32, unspliced HIV RNA aOR: 1.18; 95% CI 1.06-1.32) (Figure 4) and liver fibrosis (defect HIV DNA aOR: 1.28; 95% CI 1.08-1.52, total HIV RNA aOR: 1.12; 95% CI 1.00-1.17) (Figure 4). These associations remained significant after additionally adjusting for known risk factors (Table S8-12).

Using the Youden index, optimal cut-off values were identified for each outcome, stratifying participants into high-risk and low-risk (Table S13). Participants in the high-risk group had an increased risk for CVD after treatment initiation (aOR 3.03; 95% CI 1.76-5.48), carotid plaque (aOR 1.34; 95% CI 1.06-1.70), liver fibrosis (aOR 2.44; 95% CI 1.45-4.06), liver steatosis (aOR 1.42; 95% CI 1.05-1.91) and malignancy during follow-up (aOR 4.54 95% CI 2.07-9.90).

#### Effect of reservoir size on comorbidities was independent from systemic inflammation

Systemic inflammation may cause or mediate the effect of reservoir on comorbidities. Using plasma proteome data, we added known HIV-related inflammatory markers, including sCD14, sCD163, CXCL10, IFNγ, IL10, TNFα, IL1β and IL6 into the models [28] (Table S8-12), and found that adjusted estimates remained unchanged for total HIV DNA, defect HIV DNA, total HIV RNA and unspliced HIV RNA, suggesting there is an excess effect of the reservoir that is not explained by typical inflammatory plasma proteomic profiles.

## Discussion

In the present observational study we show that larger reservoir size (HIV DNA and HIV RNA) associates with older biological aging scores and higher comorbidities risk, such as cardiovascular diseases, liver fibrosis and cancer. We also provide evidence that this effect is not purely dependent on systemic inflammation. These findings underscore the impact of reservoir size, apart from the traditional risk factors and chronic inflammation, on health outcomes and argue for studies to explore the effect of reservoir-reducing therapies to lower the risk for morbidities. Furthermore, we identified subgroups of people with HIV who have smaller reservoir sizes such as those with early initiation of antiretroviral treatment and spontaneous controllers, whereas individuals with a low CD4 nadir, individuals experiencing non-structured treatment interruptions or delayed virological response to treatment have larger reservoir sizes. Notably, reservoir markers were highly correlated, and consistent patterns across HIV DNA and HIV RNA markers were therefore expected. Moreover, defective proviruses displayed the strongest association with HIV 1 RNA levels, supporting the idea that much of the measurable HIV transcription in ART treated individuals is derived from defective proviruses rather than intact genomes capable of producing replication-competent virus [29–31].

Reservoir-reducing therapies currently aim to eliminate the intact latent reservoir so that antiretroviral therapy can be stopped, Our data raise concerns that after elimination of the replication-competent virus, a large remaining defective reservoir may still increase the development of comorbidities. Especially at risk are those that initiate treatment late and with low CD4 nadir, that may concern 30-70% of people with HIV in certain populations [7, 8]. Particularly in these people, the impact of reservoir-reducing therapies on development of comorbidities in this population should be studied. Conversely, cure-focused interventions may benefit from focusing on individuals with a small reservoir, such as young women that start antiretroviral therapy early.

The question is how the reservoir size may impact health outcomes? It is well established that chronic immune activation and inflammation increase risk for comorbidities, also in virally suppressed people with HIV [32]. Our data suggest that there is an excess risk between reservoir size and health outcomes, independent from systemic inflammation, as represented by well-known HIV-related inflammatory markers. The effect of the reservoir is also independent from CD4 nadir which may translate in immune dysfunction, even after years of successful antiretroviral treatment [33]). It is likely that the reservoir effect is tissue specific and different for HIV DNA and HIV RNA. It is known for instance that proviral integration in host DNA is not random and may localize within host genes involved in immune signalling or cellular proliferation, thereby disrupting these pathways [17]. Additionally, a fraction of the quantified HIV DNA may exist in unintegrated forms, such as LTR circles [34], which could be sensed by innate immune DNA sensors like cyclic GMP-AMP synthase (cGAS) or IFN inducible protein 16 (IFI16) [35]. On the other hand, cell-associated HIV RNA transcripts can trigger TLR7/8 sensing, causing cellular senescence and accelerated aging within tissues [36]. Moreover, HIV RNA transcription may lead to Tat or Nef, leading to epigenetic modifications or mitochondrial stress, which may also drive aging [37].

Our study has strength and limitations. The large sample size, prospective design and reservoir measurements in large number of participants present strength. However, 2-year follow-up period limits collection of incident comorbidities. Furthermore not all participants could be classified into specific clinical phenotypes, limiting power in these analyses. In addition, while digital PCR provides sensitive and precise quantification of HIV reservoir markers, it does not fully capture the replication-competence or functional activity of the reservoir, which may limit biological interpretation. In addition, technical factors, including primer or probe mismatches and variability in cDNA synthesis efficiency, may have introduced measurement bias. Participants in this study were diagnosed during a period when HIV treatment guidelines were shifting toward initiating treatment regardless of CD4 count, which adds an additional layer of complexity. This transition likely contributed to the limited number of treatment-naïve HIV controllers and made it more difficult to assess the impact of specific antiretroviral drug classes, regimens, and treatment duration on reservoir levels. The number of included women was also limited, despite extra efforts to include them. The study participants were recruited in specialized HIV treatment centers in the Netherlands, possibly limiting generalizability to geographically different populations. Additionally, this study took place during the COVID19-pandemic which did not seem to impact proviral HIV DNA but did impact plasma protein concentrations due to limited exposure to pathogens [25]. While we performed extensive sensitivity analyses and covariate adjustment, residual confounding cannot be ruled out, however across all analysis and cohorts, there were only minor changes in estimates, suggesting robust results.

In conclusion, the HIV reservoir is significantly associated with long-term health outcomes in people with HIV using antiretroviral therapy, emphasizing the need for studies investigating the effect of reservoir reducing interventions on comorbidities.

## Acknowledgements

We are deeply grateful to all participants for their generous contribution to this study. We also thank the dedicated efforts of research teams of 2000HIV for their collaboration and support. This work was supported by ViiV Healthcare (A18-1052).

## Authors Contributions

MD, TO, WT, LJ, MN, AV and LV contributed to conceptualization of the study. MD, KM, MV and EB performed reservoir experiments. MD, TO, NV, KM, YZ analyzed the data. EB, CR, JS and MB coordinated the study. MD and TO wrote the original draft of the manuscript. VR, VM, SR, JDS, SG, JvL, WT, AV and LV provided scientific input. All authors reviewed and approved the final version of the manuscript and had final responsibility for the decision to submit for publication.

## Declaration of generative AI and AI-assisted technologies in the manuscript preparation process

During the preparation of this work the author(s) used generative AI to assist with grammar and sentence formation. After using this tool, the author(s) reviewed and edited the content as needed and take(s) full responsibility for the content of the published article.

## Supplementary Figures

**Figure S1:**
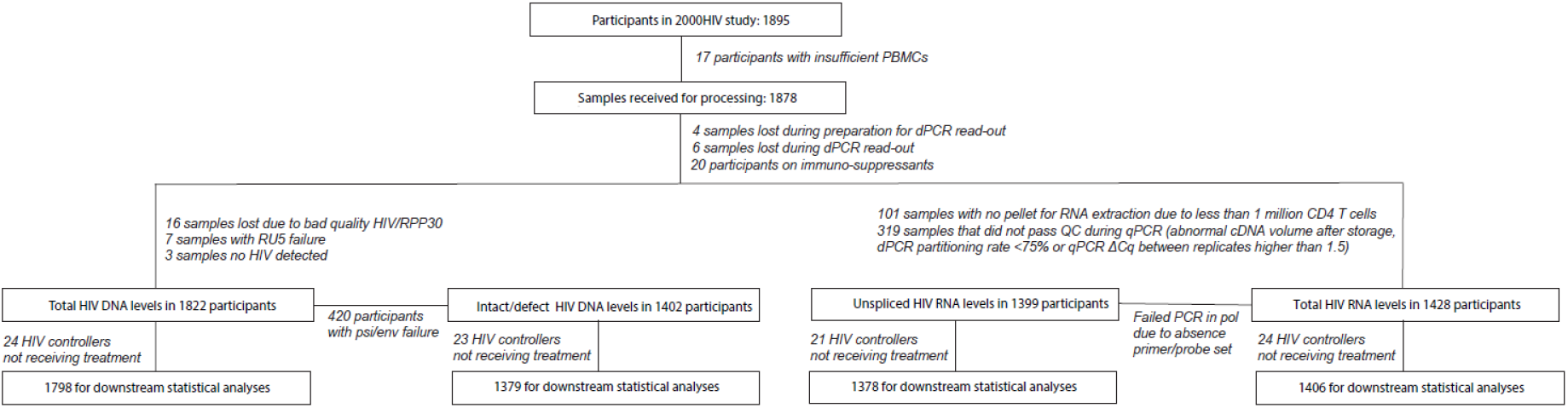
Flowchart outlining samples excluded in 2000HIV study for downstream analyses. Starting from 1895 participants, samples were excluded due to insufficient PBMCs, technical failures during dPCR processing and quality control issues. Total HIV DNA levels and intact HIV DNA were successfully measured in 1822 and 1402 participants, respectively. Total and unspliced cell- associated HIV RNA levels were successfully measured in 1428 and 1399 participant, respectively. Participants not receiving antiretroviral therapy were excluded for downstream statistical analyses.

**Figure S2:**
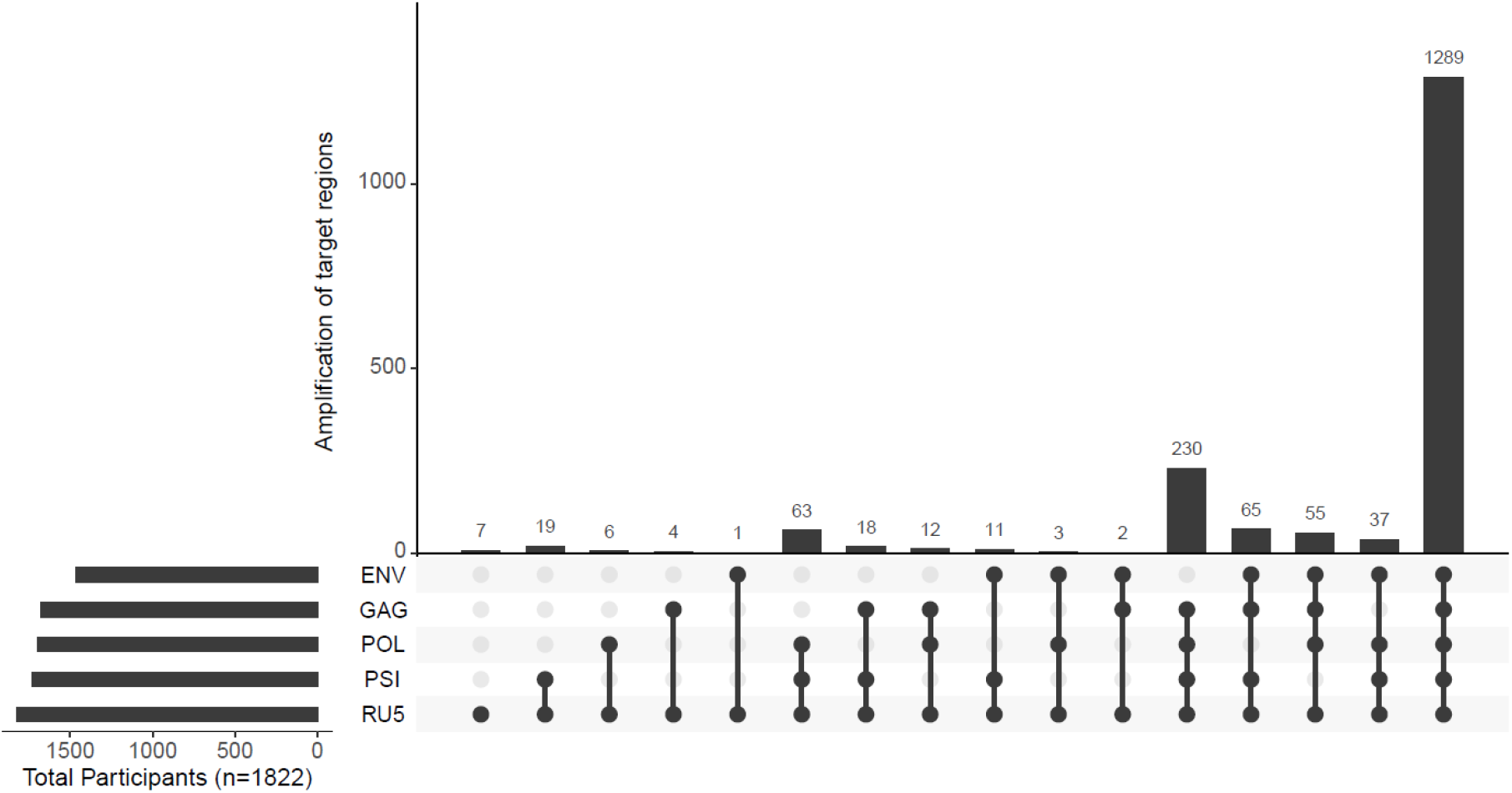
Upset plot with amplification results of 5 target regions of the Rainbow proviral HIV DNA assay in participants (n = 1822). The horizontal bars depict in how many participants each target region was successfully detected. The vertical bars demonstrate the number of participants where a combination of different target regions was successful. A grey dot means that there was a loss of signal for that region, and a black dot means that partitions were picked up above the limit of blank.

**Figure S3:**
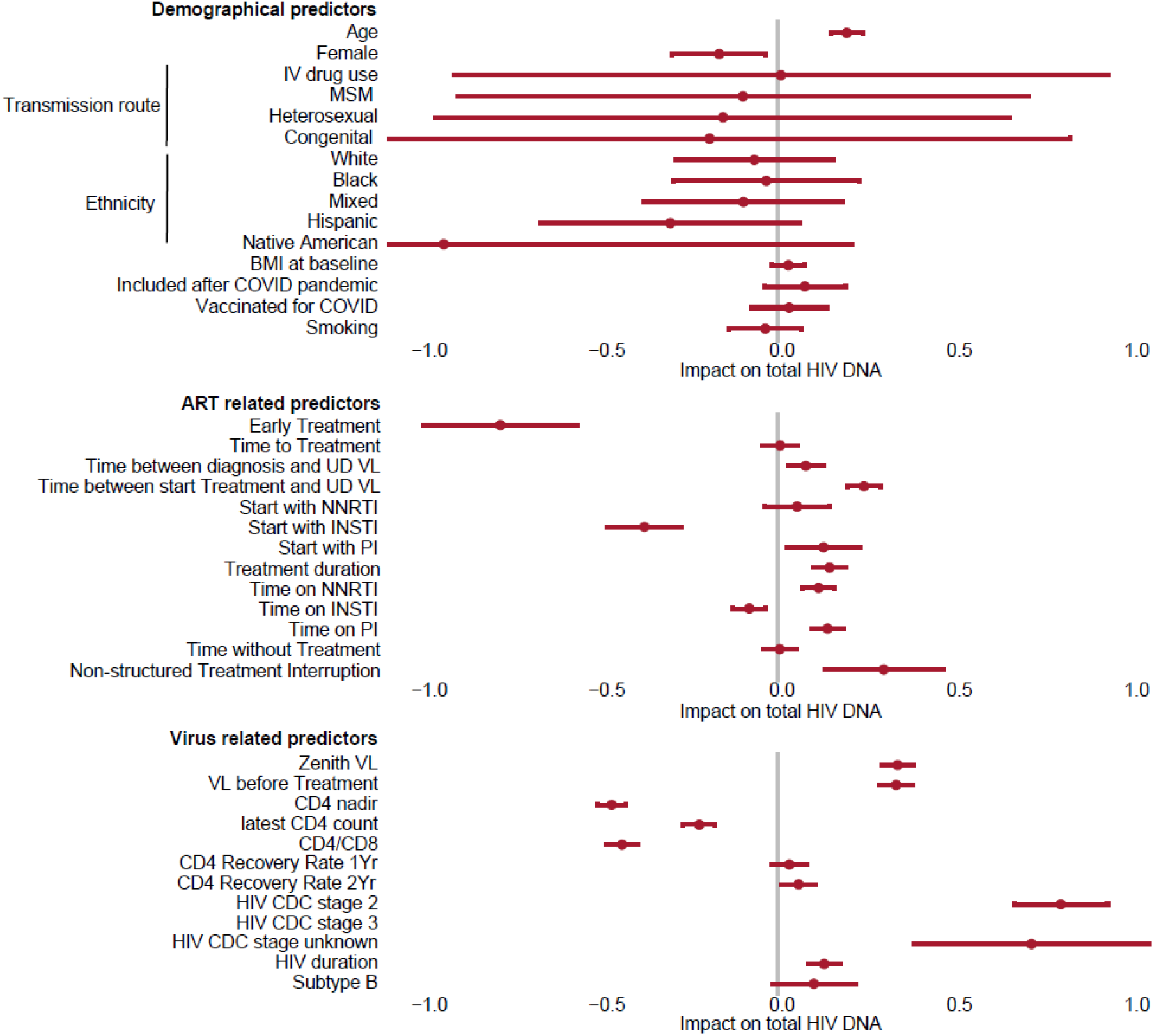
Coefficient plot of univariate analysis with total HIV as outcome variable and predictor variables on the y-axis.

**Figure S4:**
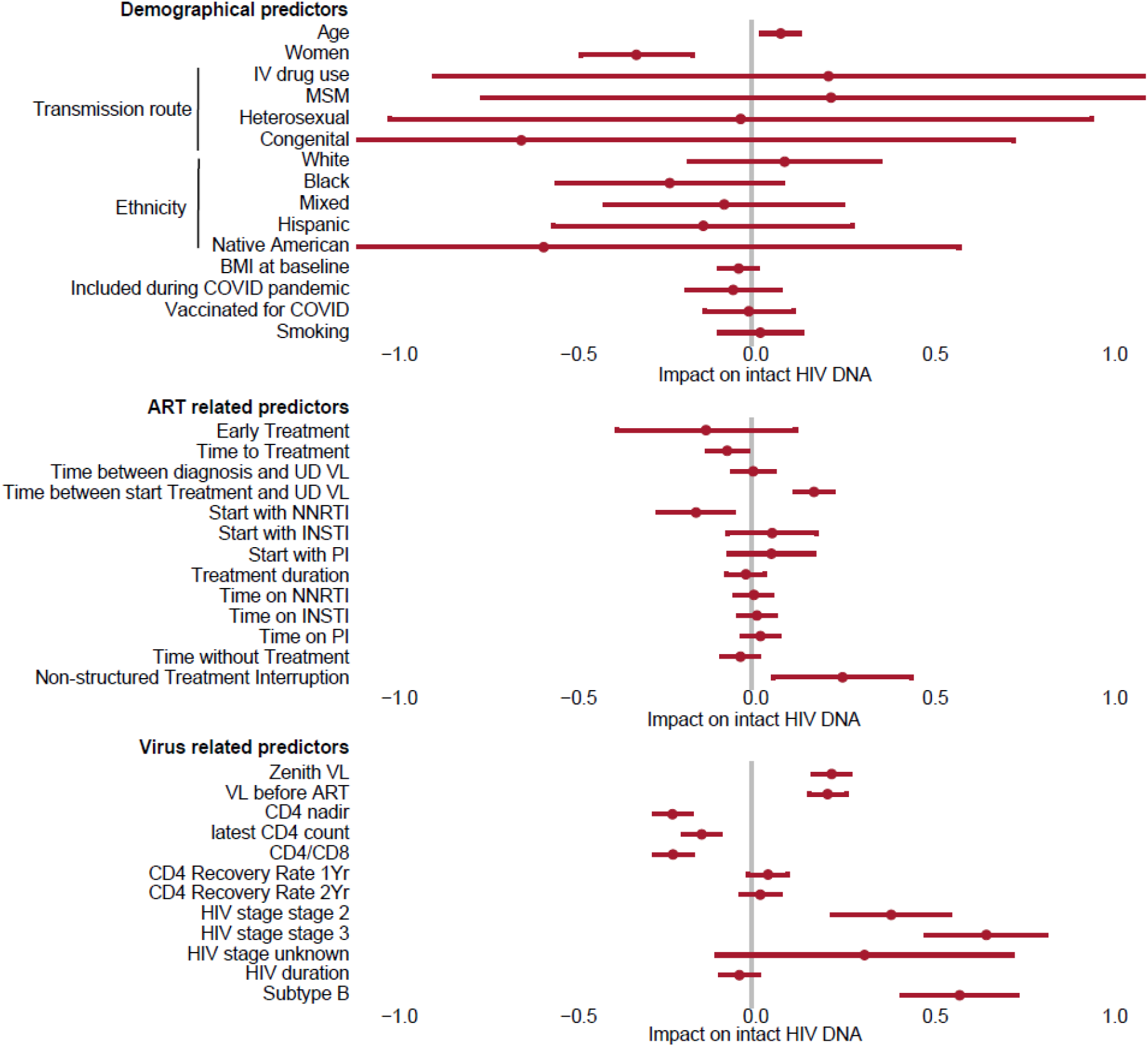
Coefficient plot of univariate analysis with intact HIV DNA levels as outcome variable and predictor variables on the y-axis.

**Figure S5:**
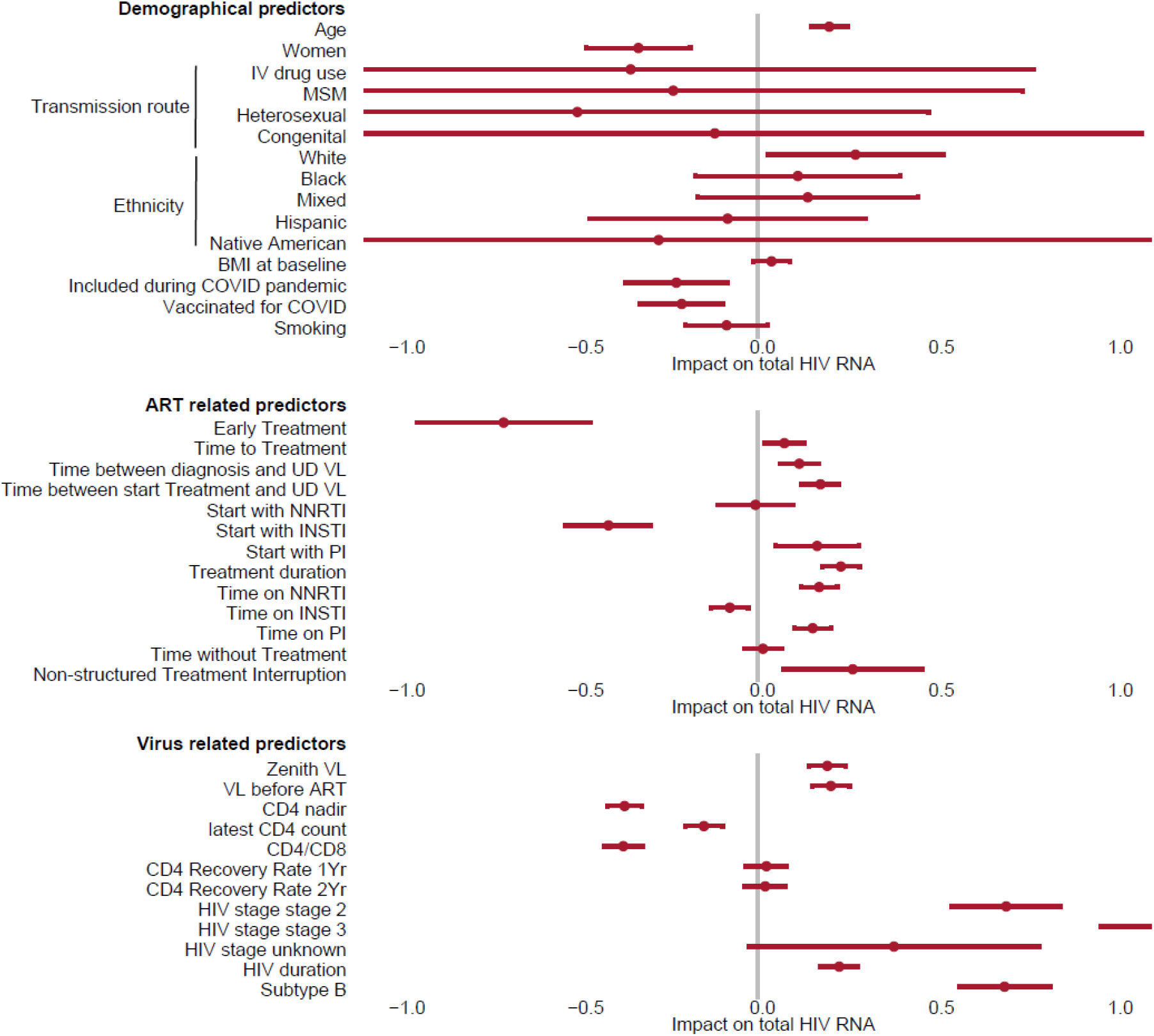
Coefficient plot of univariate analysis with total HIV RNA levels as outcome variable and predictor variables on the y-axis.

**Figure S6:**
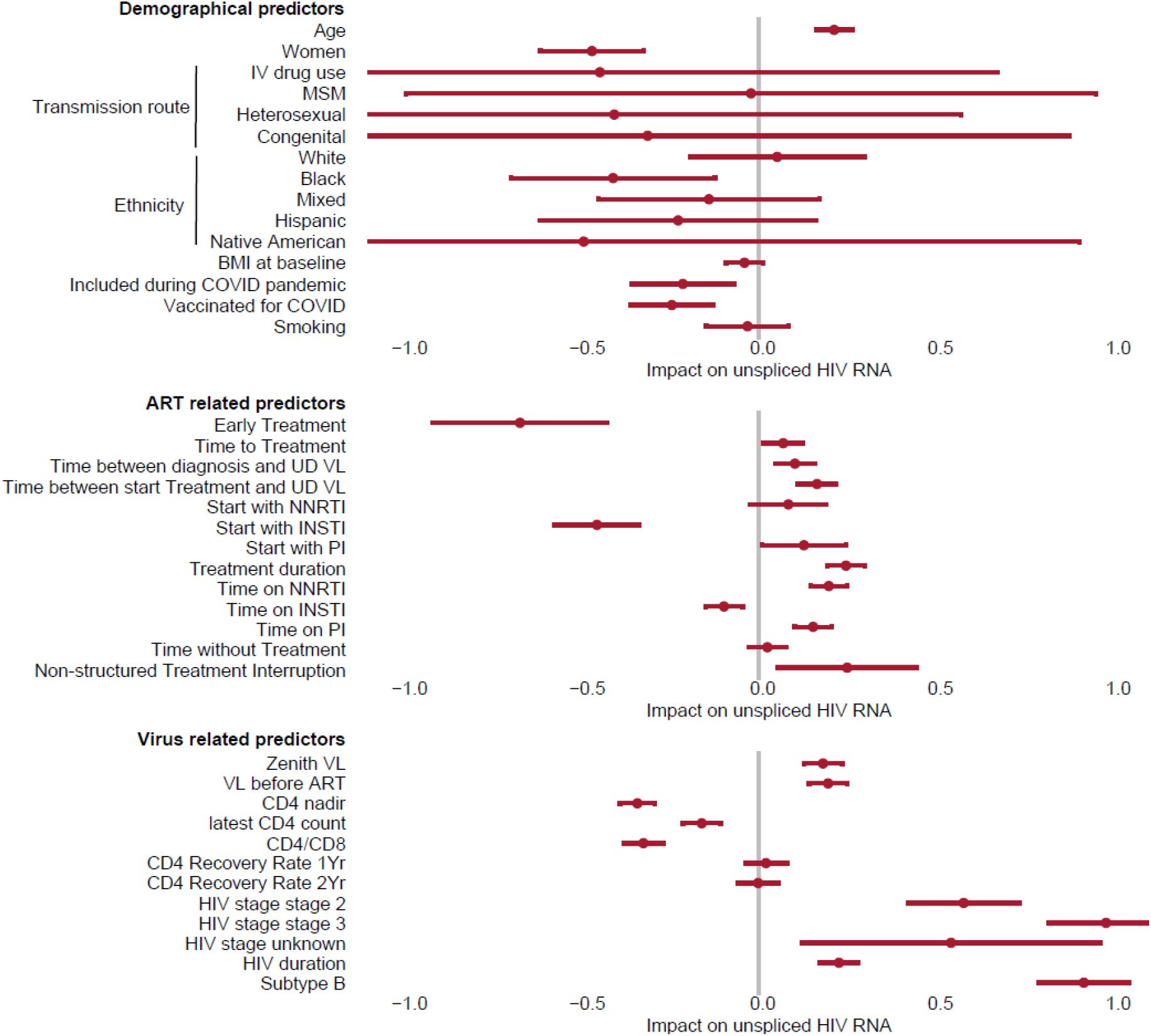
Coefficient plot of univariate analysis with unspliced HIV RNA levels as outcome variable and predictor variables on the y-axis.

**Figure S7:**
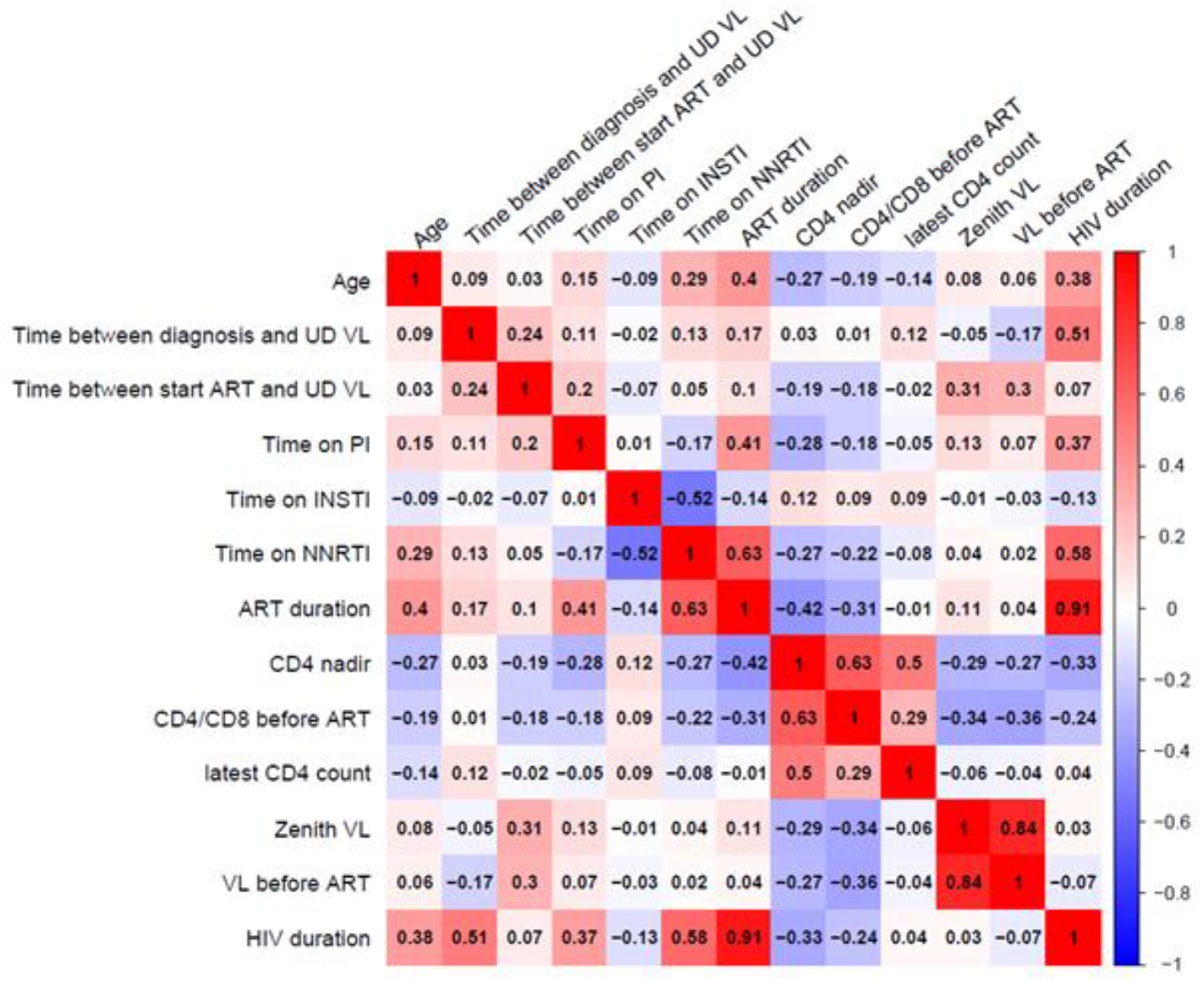
Correlation plot for continuous variables in univariate analysis.

**Figure S8:**
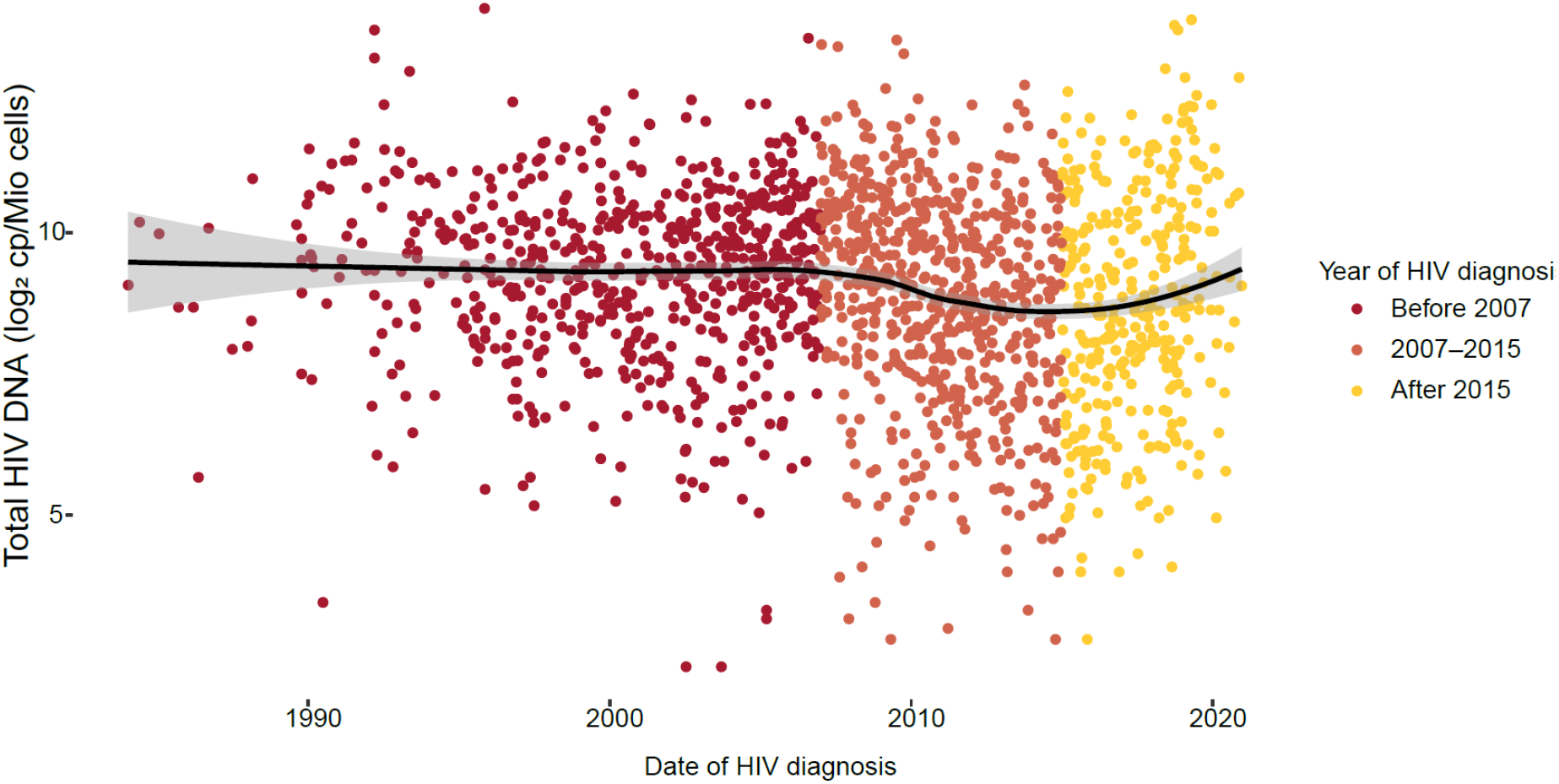
LOESS analysis of total HIV DNA levels by date of HIV diagnosis. Participants were stratified into 3 groups based on the year of their HIV diagnosis.

**Figure S9:**
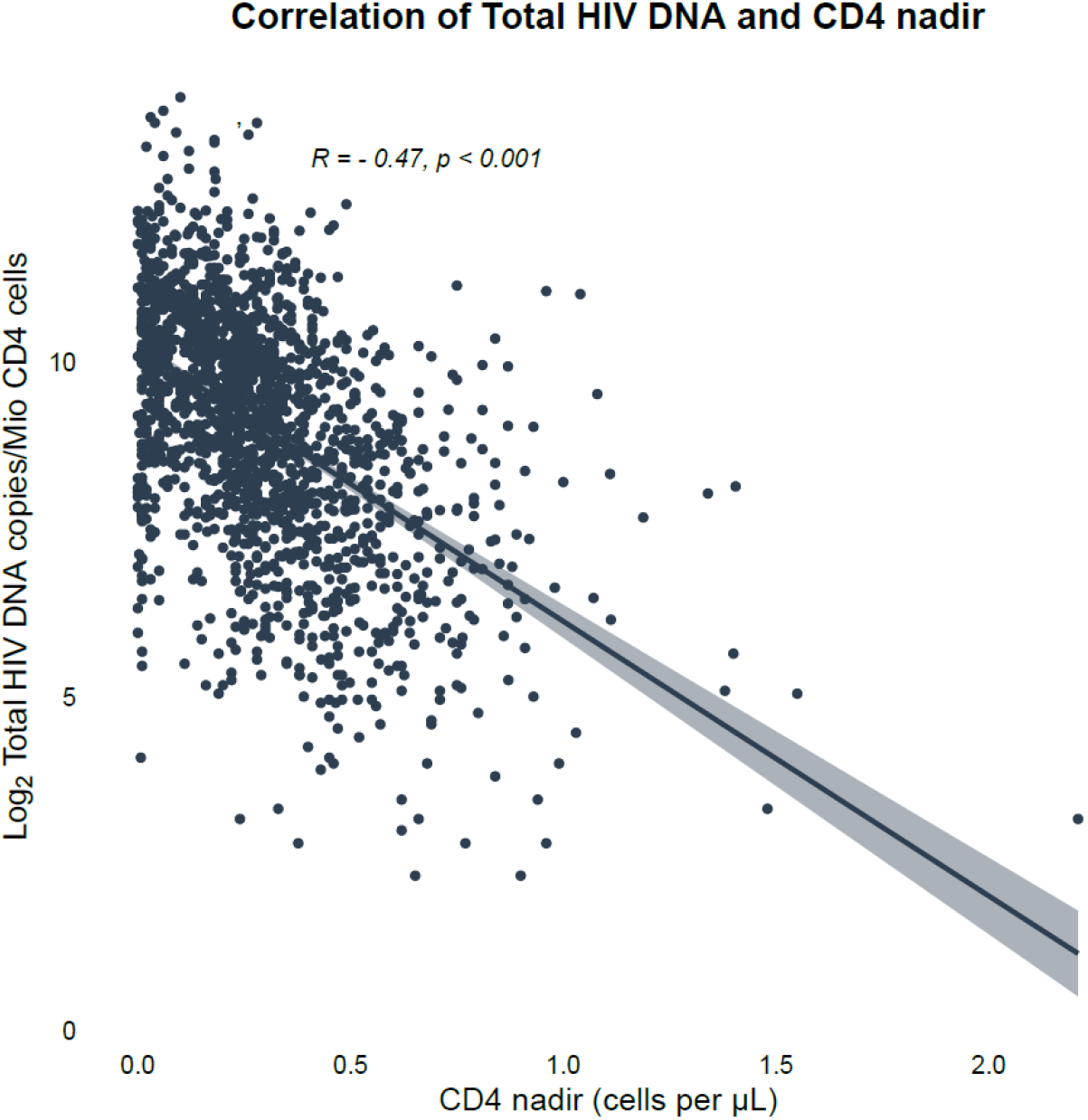
Correlation plot between CD4 nadir and total HIV DNA levels (Pearson correlation).

## Supplementary tables

**Table S1:** Operation definitions.

| **HIV phenotypes** |  |
|---|---|
| **HIV controllers** |  |
| Elite controllers (EC) | HIV-positive, ≥1 year without cART and ≥3 consecutive HIV-RNA <75 copies/mL spanning ≥12 months and stable CD4 ≥500 cells/µL. |
| Viremic controllers (VC) | HIV-positive, ≥5 years without cART and always HIV-RNA <10,000 copies/mL and stable CD4 ≥500 cells/µL |
| Transient controllers (TC) | Met EC or VC criteria at some point but initiated ART due to viral load >10,000 copies/mL or CD4 <500 cells/µL |
| Non-controllers | Eligible to meet EC or VC criteria but did not. |
| **Rapid progressors** |  |
| Yes | ≤1 year from last negative HIV test or confirmed acute HIV to CD4 <350 cells/µL, or ≤2 years to CD4 <200 cells/µL. |
| No | Not satisfying the above |
| Unknown | Start ART before rapid progressor criteria could be satisfied, CD4 < 350 cells/µL within 1 year of non-acute HIV diagnosis |
| **Immunologic response** |  |
| Immunological non-responder (INR) | CD4 <350 cells/µL after ≥2 years of suppressive cART. |
| Immunological responder (IR): | CD4 >500 cells/µL under suppressive cART. |
| Undefined | CD4 >350 but <500 cells/µL, missing data, or EC without cART. |
| CD4 recovery: | Increment in CD4 count after starting ART at 1 year (or 2 years if CD4 nadir <500). |
| **ART categories** |  |
| Early cART | Initiation <1 month after confirmed acute HIV infection (last negative HIV test <6 months prior or incomplete immunoblot at diagnosis). |
| Not early cART | Complete immunoblot at diagnosis, >180 days between HIV diagnosis and cART initiation (if exact dates available), or ≥2 years between diagnosis and cART start (if only year known). |
| Non-structured treatment interruption | Self-reported interruption >3 months after initiation |
| **Cardiovascular disease (CVD)** |  |
| Clinical CVD | Stroke, transient ischemic attack, myocardial infarction, angina pectoris, or peripheral artery disease diagnosed by a relevant specialist according to current guidelines. |
| CVD after start treatment | Only events diagnosed after treatment initiation. |
| Incident CVD during follow-up | Same definition as above; participants with prior CVD excluded. |
| **Liver morbidities trough transient elastography using FibroScan at cohort entry** |  |
| Liver steatosis | Controlled attenuation parameter (CAP) ≥263 dB/m for steatosis |
| Liver fibrosis | Liver stiffness measurement (LSM) ≥7.0 kPa for fibrosis |
| **Viremia categories** |  |
| Residual viremia (RV): | Viral load <40 copies/mL but detectable. |
| Target not detected (TND) | Undetectable viral load. |
| Quantifiable viremia in 3 years before entry | At least one detectable viral load >40 copies/mL. |
| Blips | 40-200 copies/mL. |
| Viremia | 200-1,000 copies/mL; suspected ART non-adherence was recorded when applicable |
| Failure | >1,000 copies/mL; suspected ART non-adherence was recorded when applicable |
| Quantifiable viremia follow-up | At least one detectable viral load >50 copies/mL |

**Table S2:** Baseline characteristics of the 2000HIV study with total HIV-1 DNA levels (n = 1822). Continuous variables are presented as median (interquartile range). Categorical variables are reported as number (percentage). For each variable, sample size is highlighted as some variables included missing values for certain participants. (IV = intravenous, MSM = men having sex with men, NNRT = non-nucleoside reverse-transcriptase inhibitors, INSTI = integrase strand transfer inhibitor, PI = protease inhibitor)

| **Characteristic** | **N** | **Overall**<br>**N = 1,822** |
|---|---|---|
| Age (years) | 1,822 | 53.0 (44.0-60.0) |
| Sex at birth | 1,822 |  |
| Female |  | 270 (15%) |
| Male |  | 1,552 (85%) |
| Ethnicity | 1,820 |  |
| Asian |  | 83 (4.6%) |
| Black |  | 186 (10%) |
| Hispanic |  | 46 (2.5%) |
| Mixed |  | 125 (6.9%) |
| Native American |  | 3 (0.2%) |
| White |  | 1,377 (76%) |
| HIV transmission route | 1,736 |  |
| Blood products |  | 6 (0.3%) |
| Congenital |  | 10 (0.6%) |
| Heterosexual |  | 385 (22%) |
| IV drug use |  | 20 (1.2%) |
| MSM |  | 1,315 (76%) |
| BMI baseline | 1,822 | 24.9 (22.4-27.6) |
| Collection during pandemic | 1,822 |  |
| Before |  | 366 (20%) |
| After |  | 1,456 (80%) |
| Received COVID19 vaccine before sampling | 1,822 |  |
| No |  | 1,365 (75%) |
| Yes |  | 457 (25%) |
| Currently smoking | 1,822 |  |
| Not smoking |  | 1,282 (70%) |
| Smoking |  | 540 (30%) |
| Treatment initiation | 1,322 |  |
| Started treatment during acute HIV infection |  | 86 (6.5%) |
| Started treatment during chronic HIV infection |  | 1,236 (93%) |
| Time to treatment (days) | 1,614 | 126.0 (31.0-853.0) |
| Time between diagnosis and undetectable viral load (years) | 1,522 | 1.0 (0.5-3.0) |
| Time between start treatment and undetectable viral load (years) | 1,641 | 0.3 (0.2-0.6) |
| First treatment with | 1,820 |  |
| NNRTI |  | 811 (45%) |
| INSTI |  | 449 (25%) |
| PI |  | 503 (28%) |
| Treatment duration | 1,797 | 10.2 (6.0-16.4) |
| Non-structured treatment interruption | 1,803 | 152 (8.4%) |
| Viral load zenith (copies/mL) | 1,659 | 100,000.0 (39,226.0-290,000.0) |
| Viral load before treatment (copies/mL) | 1,649 | 80,700.0 (24,554.0-200,000.0) |
| CD4 nadir (10^9 cells/L) | 1,788 | 0.3 (0.1-0.4) |
| CD4/CD8 ratio before start treatment | 1,474 | 0.3 (0.2-0.5) |
| Latest CD4 count (10^9 cells/L) | 1,808 | 0.7 (0.5-0.9) |
| CD4 Recovery Rate 1 Yr | 1,519 | 0.2 (0.1-0.3) |
| CD4 Recovery Rate 2Yr | 1,549 | 0.1 (0.1-0.2) |
| HIV CDC stage | 1,804 |  |
| Stage 1 (≥500 CD4 T cells/µL) |  | 237 (13%) |
| Stage 2 (200-499 CD4 T cells/µL) |  | 833 (46%) |
| Stage 3 (<200 CD4 T cells/µL or AIDS defining diagnosis) |  | 702 (39%) |
| Unknown |  | 32 (1.8%) |
| HIV duration | 1,822 | 12.5 (7.4-19.2) |
| Subtype | 1,778 |  |
| B |  | 1,438 (81%) |
| nonB |  | 340 (19%) |

**Table S3:** Results from univariable logistic regression models with total HIV DNA levels as predictor.

| outcome | OR | CI95-low | CI95-high | P value |
|---|---|---|---|---|
| AIDS defining infection | 1.389 | 1.271 | 1.522 | <0.001 |
| AIDS defining malignancy | 1.327 | 1.174 | 1.506 | <0.001 |
| Anxiety | 1.024 | 0.889 | 1.187 | 0.743 |
| Cardiovascular disease (CVD) after start treatment | 1.427 | 1.236 | 1.656 | <0.001 |
| Carotid plaque | 1.185 | 1.120 | 1.254 | <0.001 |
| COPD | 1.168 | 0.998 | 1.375 | 0.058 |
| CVD during follow-up | 1.063 | 0.877 | 1.301 | 0.541 |
| Depression | 0.999 | 0.930 | 1.074 | 0.986 |
| Diabetes melitus type 2 | 1.195 | 1.048 | 1.370 | 0.009 |
| HIV encephalopathy, PML, wasting syndrome | 1.419 | 1.272 | 1.590 | <0.001 |
| Hypercholesterolemia | 1.103 | 1.038 | 1.174 | 0.002 |
| Hypertriglyceridemia | 1.448 | 1.077 | 1.983 | 0.018 |
| Immunological non-responders | 1.506 | 1.303 | 1.751 | <0.001 |
| Inflammatory Arthritis | 0.918 | 0.745 | 1.143 | 0.432 |
| Irritable bowel disease | 0.994 | 0.766 | 1.312 | 0.967 |
| Liver fibrosis | 1.227 | 1.070 | 1.413 | 0.004 |
| Liver steatosis | 1.135 | 1.049 | 1.229 | 0.002 |
| Liver steatosis during follow-up | 0.898 | 0.778 | 1.040 | 0.148 |
| Non-AIDS defining malignancy | 1.073 | 0.943 | 1.226 | 0.294 |
| Non-AIDS Defining Malignancy during follow-up | 1.272 | 1.026 | 1.596 | 0.033 |
| Osteoporosis | 1.056 | 0.908 | 1.234 | 0.489 |
| Pulmonary disease | 1.037 | 0.955 | 1.127 | 0.397 |
| Quantifiable viremia 3 years before study entry | 1.391 | 1.286 | 1.507 | <0.001 |
| Quantifiable viremia during follow-up | 1.465 | 1.299 | 1.660 | <0.001 |
| Residual viremia 3 years before study entry | 1.313 | 1.234 | 1.399 | <0.001 |
| Residual viremia during follow-up | 1.559 | 1.412 | 1.729 | <0.001 |

**Table S4:** Results from univariable logistic regression models with intact HIV DNA levels as predictor.

| outcome | OR | CI95-low | CI95-high | P value |
|---|---|---|---|---|
| AIDS defining infection | 1.068 | 0.996 | 1.143 | 0.062 |
| AIDS defining malignancy | 0.969 | 0.874 | 1.070 | 0.539 |
| Anxiety | 1.115 | 0.991 | 1.252 | 0.067 |
| Cardiovascular disease (CVD) after start treatment | 1.142 | 1.021 | 1.275 | 0.019 |
| Carotid plaque | 1.032 | 0.983 | 1.084 | 0.2 |
| COPD | 1.143 | 1.006 | 1.295 | 0.039 |
| CVD during follow-up | 1.042 | 0.874 | 1.232 | 0.64 |
| Depression | 1.030 | 0.967 | 1.096 | 0.36 |
| Diabetes melitus type 2 | 1.030 | 0.925 | 1.145 | 0.581 |
| HIV encephalopathy, PML, wasting syndrome | 1.109 | 1.019 | 1.207 | 0.016 |
| Hypercholesterolemia | 1.039 | 0.986 | 1.095 | 0.149 |
| Hypertriglyceridemia | 1.141 | 0.879 | 1.465 | 0.306 |
| Immunological non-responders | 1.206 | 1.080 | 1.347 | 0.001 |
| Inflammatory Arthritis | 0.872 | 0.703 | 1.061 | 0.19 |
| Irritable bowel disease | 1.178 | 0.945 | 1.459 | 0.137 |
| Liver fibrosis | 1.102 | 0.991 | 1.224 | 0.07 |
| Liver steatosis | 0.992 | 0.927 | 1.060 | 0.803 |
| Liver steatosis during follow-up | 0.950 | 0.833 | 1.078 | 0.439 |
| Non-AIDS defining malignancy | 1.013 | 0.904 | 1.130 | 0.826 |
| Non-AIDS Defining Malignancy during follow-up | 1.192 | 0.995 | 1.424 | 0.054 |
| Osteoporosis | 1.072 | 0.941 | 1.217 | 0.286 |
| Pulmonary disease | 1.024 | 0.952 | 1.100 | 0.517 |
| Quantifiable viremia 3 years before study entry | 1.158 | 1.089 | 1.231 | <0.001 |
| Quantifiable viremia during follow-up | 1.156 | 1.054 | 1.267 | 0.002 |
| Residual viremia 3 years before study entry | 1.170 | 1.111 | 1.233 | <0.001 |
| Residual viremia during follow-up | 1.311 | 1.213 | 1.421 | <0.001 |

**Table S5:**
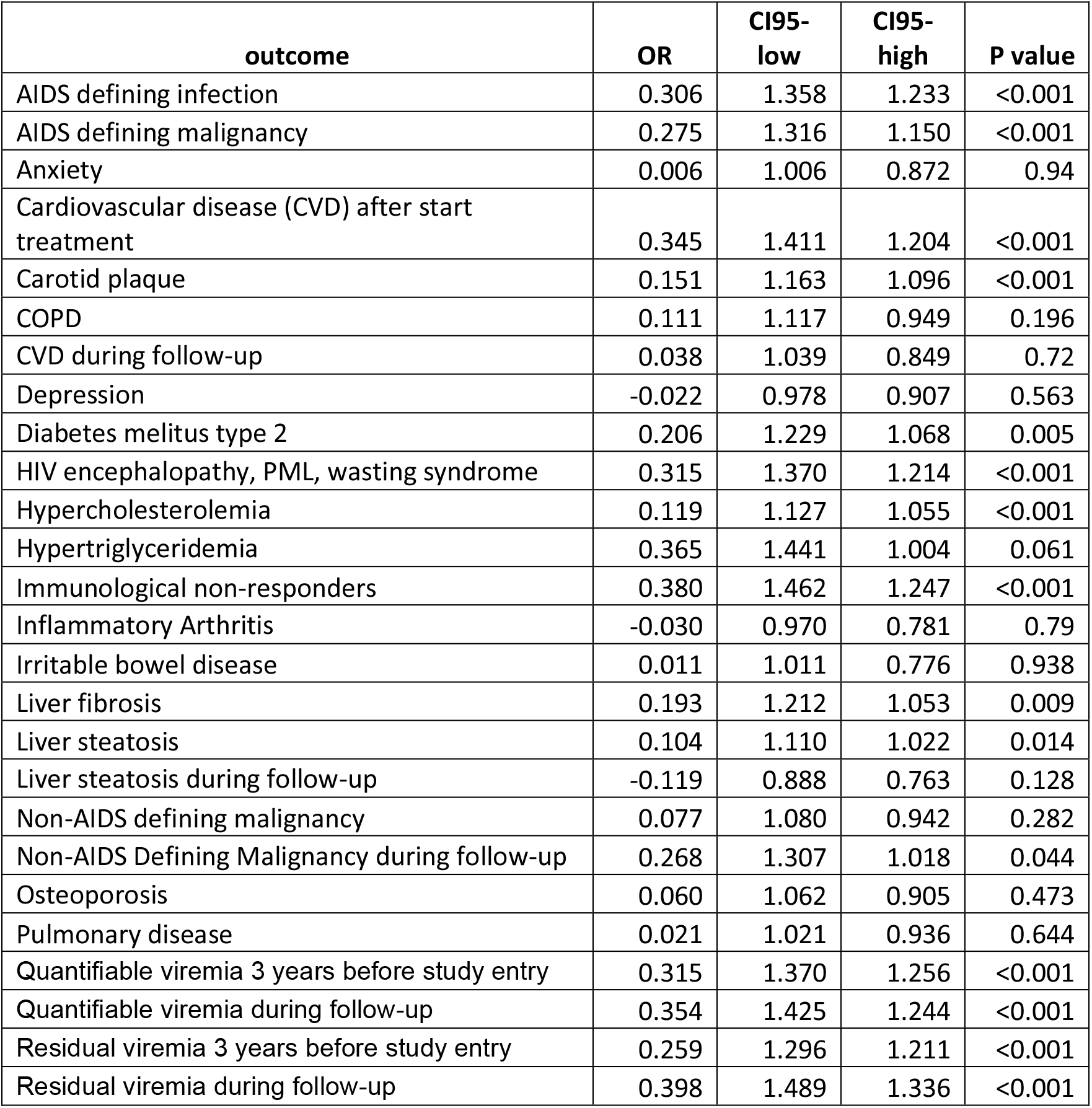
Results from univariable logistic regression models with defect HIV DNA levels as predictor.

**Table S6:** Results from univariable logistic regression models with total HIV RNA levels as predictor.

| **outcome** | **OR** | **CI95-low** | **CI95-high** | **P value** |
|---|---|---|---|---|
| AIDS defining infection | 0.170 | 1.186 | 1.112 | <0.001 |
| AIDS defining malignancy | 0.202 | 1.223 | 1.114 | <0.001 |
| Anxiety | 0.009 | 1.009 | 0.917 | 0.863 |
| Cardiovascular disease (CVD) after start treatment | 0.183 | 1.200 | 1.083 | 0.001 |
| Carotid plaque | 0.071 | 1.074 | 1.032 | <0.001 |
| COPD | 0.124 | 1.132 | 0.997 | 0.062 |
| CVD during follow-up | -0.082 | 0.921 | 0.798 | 0.271 |
| Depression | 0.015 | 1.015 | 0.964 | 0.568 |
| Diabetes melitus type 2 | 0.152 | 1.164 | 1.057 | 0.002 |
| HIV encephalopathy, PML, wasting syndrome | 0.190 | 1.210 | 1.118 | <0.001 |
| Hypercholesterolemia | 0.096 | 1.101 | 1.052 | <0.001 |
| Hypertriglyceridemia | 0.337 | 1.401 | 1.118 | 0.005 |
| Immunological non-responders | 0.173 | 1.189 | 1.060 | 0.004 |
| Inflammatory Arthritis | 0.048 | 1.049 | 0.899 | 0.554 |
| Irritable bowel disease | -0.092 | 0.912 | 0.731 | 0.428 |
| Liver steatosis | 0.086 | 1.090 | 1.030 | 0.003 |
| Liver steatosis during follow-up | 0.046 | 1.047 | 0.932 | 0.449 |
| liver_fibrosis_new_excl | 0.103 | 1.109 | 1.010 | 0.033 |
| Non-AIDS defining malignancy | 0.119 | 1.127 | 1.024 | 0.017 |
| Non-AIDS Defining Malignancy during follow-up | 0.141 | 1.151 | 0.987 | 0.083 |
| Osteoporosis | 0.064 | 1.066 | 0.954 | 0.272 |
| Pulmonary disease | 0.024 | 1.025 | 0.965 | 0.431 |
| Quantifiable viremia 3 years before study entry | 0.125 | 1.133 | 1.076 | <0.001 |
| Quantifiable viremia during follow-up | 0.169 | 1.184 | 1.089 | <0.001 |
| Residual viremia 3 years before study entry | 0.115 | 1.122 | 1.074 | <0.001 |
| Residual viremia during follow-up | 0.175 | 1.191 | 1.115 | <0.001 |

**Table S7:** Results from univariable logistic regression models with unspliced HIV RNA levels as predictor.

| outcome | OR | CI95-low | CI95-high | P value |
|---|---|---|---|---|
| AIDS defining infection | 1.171 | 1.105 | 1.243 | <0.001 |
| AIDS defining malignancy | 1.196 | 1.099 | 1.304 | <0.001. |
| Anxiety | 0.965 | 0.875 | 1.063 | 0.472 |
| Cardiovascular disease (CVD) after start treatment | 1.205 | 1.097 | 1.327 | <0.001 |
| Carotid plaque | 1.077 | 1.036 | 1.120 | <0.001 |
| COPD | 1.152 | 1.018 | 1.306 | 0.026 |
| CVD during follow-up | 0.987 | 0.845 | 1.148 | 0.867 |
| Depression | 1.011 | 0.961 | 1.064 | 0.674 |
| Diabetes melitus type 2 | 1.130 | 1.035 | 1.235 | 0.007 |
| HIV encephalopathy, PML, wasting syndrome | 1.202 | 1.120 | 1.292 | <0.001 |
| Hypercholesterolemia | 1.087 | 1.041 | 1.136 | <0.001 |
| Hypertriglyceridemia | 1.265 | 1.046 | 1.541 | 0.017 |
| Immunological non-responders | 1.226 | 1.104 | 1.365 | <0.001 |
| Inflammatory Arthritis | 1.096 | 0.939 | 1.280 | 0.245 |
| Irritable bowel disease | 0.939 | 0.730 | 1.190 | 0.61 |
| Liver fibrosis | 1.023 | 0.938 | 1.115 | 0.604 |
| Liver steatosis | 1.058 | 1.002 | 1.117 | 0.042 |
| Liver steatosis during follow-up | 1.031 | 0.923 | 1.151 | 0.589 |
| Non-AIDS defining malignancy | 1.175 | 1.074 | 1.287 | <0.001 |
| Non-AIDS Defining Malignancy during follow-up | 1.136 | 0.986 | 1.311 | 0.079 |
| Osteoporosis | 1.108 | 0.995 | 1.235 | 0.062 |
| Pulmonary disease | 1.052 | 0.992 | 1.117 | 0.091 |
| Quantifiable viremia 3 years before study entry | 1.121 | 1.068 | 1.178 | <0.001 |
| Quantifiable viremia during follow-up | 1.094 | 1.015 | 1.179 | 0.019 |
| Residual viremia 3 years before study entry | 1.080 | 1.037 | 1.125 | <0.001 |
| Residual viremia during follow-up | 1.101 | 1.038 | 1.169 | 0.002 |

**Table S8:** Univariable and mutlivariable models for different outcomes with total HIV DNA as predictor. (N = sample size data in model, Cases = events of outcome, OR = odds ratio, CI_Low = lower limit 95%CI, CI_Up = upper limit 95%CI)

| Outcome | N | Cases | OR | CI_Low | CI_Up | p.value | Adjusted for: |
|---|---|---|---|---|---|---|---|
| Carotid plaque | 1721 | 870 | 1.185 | 1.120 | 1.254 | <0.001 | / |
| Carotid plaque | 1688 | 854 | 1.109 | 1.034 | 1.191 | 0.004 | Age, sex, CD4 nadir |
| Carotid plaque | 1535 | 757 | 1.115 | 1.035 | 1.203 | 0.005 | Age, sex, CD4 nadir, BMI, HT, HC, Fa Hx, DM, current smoking, time to treatment and treatment duration |
| Carotid plaque | 1520 | 748 | 1.120 | 1.038 | 1.209 | 0.004 | Age, sex, CD4 nadir, BMI, HT, HC, Fa Hx, DM, current smoking, time to treatment and treatment duration, IL1B, TNF, IL6, CXCL10, CD14, CD163, IFNG |
| CVD after treatment initiation | 1713 | 82 | 1.427 | 1.236 | 1.656 | <0.001 | / |
| CVD after treatment initiation | 1679 | 81 | 1.420 | 1.203 | 1.685 | <0.001 | Age, sex, CD4 nadir |
| CVD after treatment initiation | 1521 | 65 | 1.595 | 1.300 | 1.976 | <0.001 | Age, sex, CD4 nadir, BMI, HT, HC, Fa Hx, DM, current smoking, time to treatment and treatment duration |
| CVD after treatment initiation | 1507 | 64 | 1.595 | 1.293 | 1.989 | <0.001 | Age, sex, CD4 nadir, BMI, HT, HC, Fa Hx, DM, current smoking, time to treatment and treatment duration, IL1B, TNF, IL6, CXCL10, CD14, CD163, IFNG |
| Liver fibrosis | 930 | 84 | 1.227 | 1.070 | 1.413 | 0.004 | / |
| Liver fibrosis | 910 | 81 | 1.284 | 1.093 | 1.515 | 0.003 | Age, sex, CD4 nadir |
| Liver fibrosis | 842 | 72 | 1.241 | 1.041 | 1.487 | 0.018 | Age, sex, CD4 nadir, BMI, time to treatment and treatment duration |
| Liver fibrosis | 835 | 71 | 1.203 | 1.009 | 1.443 | 0.042 | Age, sex, CD4 nadir, BMI, time to treatment and treatment duration, IL1B, TNF, IL6, CXCL10, CD14, CD163, IFNG |
| Malignancy during follow-up | 1782 | 32 | 1.272 | 1.026 | 1.596 | 0.033 | / |
| Malignancy during follow-up | 1748 | 32 | 1.347 | 1.052 | 1.740 | 0.020 | Age, sex, CD4 nadir |
| Malignancy during follow-up | 1587 | 27 | 1.332 | 1.019 | 1.760 | 0.040 | Age, sex, CD4 nadir, history of malignancies, time to treatment and treatment duration |
| Malignancy during follow-up | 1572 | 27 | 1.316 | 1.005 | 1.753 | 0.053 | Age, sex, CD4 nadir, history of malignancies, time to treatment and treatment duration, IL1B, TNF, IL6, CXCL10, CD14, CD163, IFNG |
| Quantifiable viremia 3 yrs before study entry | 1788 | 335 | 1.391 | 1.286 | 1.507 | <0.001 | / |
| Quantifiable viremia 3 yrs before study entry | 1756 | 326 | 1.449 | 1.325 | 1.589 | <0.001 | Age, sex, CD4 nadir |
| Quantifiable viremia 3 yrs before study entry | 1595 | 303 | 1.439 | 1.309 | 1.585 | <0.001 | Age, sex, CD4 nadir, time to treatment and treatment duration |
| Quantifiable viremia 3 yrs before study entry | 1580 | 298 | 1.434 | 1.305 | 1.581 | <0.001 | Age, sex, CD4 nadir, time to treatment and treatment duration, IL1B, TNF, IL6, CXCL10, CD14, CD163, IFNG |
| Quantifiable viremia during follow-up | 1779 | 124 | 1.465 | 1.299 | 1.660 | <0.001 | / |
| Quantifiable viremia during follow-up | 1745 | 122 | 1.524 | 1.331 | 1.753 | <0.001 | Age, sex, CD4 nadir |
| Quantifiable viremia during follow-up | 1586 | 109 | 1.540 | 1.330 | 1.790 | <0.001 | Age, sex, CD4 nadir, time to treatment and treatment duration |
| Quantifiable viremia during follow-up | 1571 | 109 | 1.515 | 1.309 | 1.761 | <0.001 | Age, sex, CD4 nadir, time to treatment and treatment duration, IL1B, TNF, IL6, CXCL10, CD14, CD163, IFNG |
| Residual viremia at study entry | 1746 | 631 | 1.313 | 1.234 | 1.399 | <0.001 | / |
| Residual viremia at study entry | 1715 | 618 | 1.387 | 1.290 | 1.493 | <0.001 | Age, sex, CD4 nadir |
| Residual viremia at study entry | 1561 | 563 | 1.368 | 1.267 | 1.480 | <0.001 | Age, sex, CD4 nadir, time to treatment and treatment duration |
| Residual viremia at study entry | 1546 | 558 | 1.368 | 1.266 | 1.482 | <0.001 | Age, sex, CD4 nadir, time to treatment and treatment duration, IL1B, TNF, IL6, CXCL10, CD14, CD163, IFNG |
| Residual viremia during follow-up | 924 | 264 | 1.559 | 1.412 | 1.729 | <0.001 | / |
| Residual viremia during follow-up | 907 | 260 | 1.733 | 1.541 | 1.960 | <0.001 | Age, sex, CD4 nadir |
| Residual viremia during follow-up | 809 | 232 | 1.736 | 1.530 | 1.982 | <0.001 | Age, sex, CD4 nadir, time to treatment and treatment duration |
| Residual viremia during follow-up | 803 | 230 | 1.731 | 1.524 | 1.978 | <0.001 | Age, sex, CD4 nadir, time to treatment and treatment duration, IL1B, TNF, IL6, CXCL10, CD14, CD163, IFNG |

**Table S9:** Univariable and mutlivariable models for different outcomes with intact HIV DNA as predictor. (N = sample size data in model, Cases = events of outcome, OR = odds ratio, CI_Low = lower limit 95%CI, CI_Up = upper limit 95%CI)

| Outcome | N | Cases | OR | CI_Low | CI_Up | p.value | Adjusted for: |
|---|---|---|---|---|---|---|---|
| Carotid plaque | 1321 | 684 | 1.032 | 0.983 | 1.084 | 0.200 | / |
| Carotid plaque | 1295 | 670 | 0.998 | 0.943 | 1.056 | 0.941 | Age, sex, CD4 nadir |
| Carotid plaque | 1171 | 592 | 0.993 | 0.936 | 1.053 | 0.804 | Age, sex, CD4 nadir, BMI, HT, HC, Fa Hx, DM, current smoking, time to treatment and treatment duration |
| Carotid plaque | 1158 | 584 | 0.999 | 0.941 | 1.060 | 0.964 | Age, sex, CD4 nadir, BMI, HT, HC, Fa Hx, DM, current smoking, time to treatment and treatment duration, IL1B, TNF, IL6, CXCL10, CD14, CD163, IFNG |
| CVD after treatment initiation | 1316 | 62 | 1.142 | 1.021 | 1.275 | 0.019 | / |
| CVD after treatment initiation | 1289 | 61 | 1.140 | 1.009 | 1.287 | 0.034 | Age, sex, CD4 nadir |
| CVD after treatment initiation | 1162 | 48 | 1.155 | 0.999 | 1.336 | 0.052 | Age, sex, CD4 nadir, BMI, HT, HC, Fa Hx, DM, current smoking, time to treatment and treatment duration |
| CVD after treatment initiation | 1150 | 47 | 1.166 | 1.005 | 1.357 | 0.044 | Age, sex, CD4 nadir, BMI, HT, HC, Fa Hx, DM, current smoking, time to treatment and treatment duration, IL1B, TNF, IL6, CXCL10, CD14, CD163, IFNG |
| Liver fibrosis | 724 | 71 | 1.102 | 0.991 | 1.224 | 0.070 | / |
| Liver fibrosis | 708 | 69 | 1.114 | 0.998 | 1.242 | 0.053 | Age, sex, CD4 nadir |
| Liver fibrosis | 651 | 60 | 1.099 | 0.975 | 1.238 | 0.121 | Age, sex, CD4 nadir, BMI, time to treatment and treatment duration |
| Liver fibrosis | 646 | 59 | 1.088 | 0.964 | 1.227 | 0.169 | Age, sex, CD4 nadir, BMI, time to treatment and treatment duration, IL1B, TNF, IL6, CXCL10, CD14, CD163, IFNG |
| Malignancy during follow-up | 1367 | 22 | 1.192 | 0.995 | 1.424 | 0.054 | / |
| Malignancy during follow-up | 1340 | 22 | 1.195 | 0.989 | 1.436 | 0.060 | Age, sex, CD4 nadir |
| Malignancy during follow-up | 1211 | 18 | 1.196 | 0.975 | 1.465 | 0.082 | Age, sex, CD4 nadir, history of malignancies, time to treatment and treatment duration |
| Malignancy during follow-up | 1198 | 18 | 1.187 | 0.965 | 1.462 | 0.102 | Age, sex, CD4 nadir, history of malignancies, time to treatment and treatment duration, IL1B, TNF, IL6, CXCL10, CD14, CD163, IFNG |
| Quantifiable viremia 3 yrs before study entry | 1374 | 247 | 1.158 | 1.089 | 1.231 | <0.001 | / |
| Quantifiable viremia 3 yrs before study entry | 1348 | 238 | 1.145 | 1.075 | 1.220 | <0.001 | Age, sex, CD4 nadir |
| Quantifiable viremia 3 yrs before study entry | 1218 | 222 | 1.138 | 1.065 | 1.216 | <0.001 | Age, sex, CD4 nadir, time to treatment and treatment duration |
| Quantifiable viremia 3 yrs before study entry | 1205 | 218 | 1.142 | 1.069 | 1.221 | <0.001 | Age, sex, CD4 nadir, time to treatment and treatment duration, IL1B, TNF, IL6, CXCL10, CD14, CD163, IFNG |
| Quantifiable viremia during follow-up | 1366 | 90 | 1.156 | 1.054 | 1.267 | 0.002 | / |
| Quantifiable viremia during follow-up | 1339 | 88 | 1.155 | 1.049 | 1.270 | 0.003 | Age, sex, CD4 nadir |
| Quantifiable viremia during follow-up | 1210 | 79 | 1.167 | 1.054 | 1.292 | 0.003 | Age, sex, CD4 nadir, time to treatment and treatment duration |
| Quantifiable viremia during follow-up | 1197 | 79 | 1.159 | 1.049 | 1.281 | 0.004 | Age, sex, CD4 nadir, time to treatment and treatment duration, IL1B, TNF, IL6, CXCL10, CD14, CD163, IFNG |
| Residual viremia at study entry | 1342 | 462 | 1.170 | 1.111 | 1.233 | <0.001 | / |
| Residual viremia at study entry | 1318 | 452 | 1.165 | 1.104 | 1.230 | <0.001 | Age, sex, CD4 nadir |
| Residual viremia at study entry | 1192 | 408 | 1.154 | 1.090 | 1.222 | <0.001 | Age, sex, CD4 nadir, time to treatment and treatment duration |
| Residual viremia at study entry | 1179 | 404 | 1.156 | 1.091 | 1.225 | <0.001 | Age, sex, CD4 nadir, time to treatment and treatment duration, IL1B, TNF, IL6, CXCL10, CD14, CD163, IFNG |
| Residual viremia during follow-up | 706 | 191 | 1.311 | 1.213 | 1.421 | <0.001 | / |
| Residual viremia during follow-up | 693 | 189 | 1.320 | 1.216 | 1.436 | <0.001 | Age, sex, CD4 nadir |
| Residual viremia during follow-up | 615 | 170 | 1.284 | 1.177 | 1.404 | <0.001 | Age, sex, CD4 nadir, time to treatment and treatment duration |
| Residual viremia during follow-up | 609 | 168 | 1.291 | 1.181 | 1.416 | <0.001 | Age, sex, CD4 nadir, time to treatment and treatment duration, IL1B, TNF, IL6, CXCL10, CD14, CD163, IFNG |

**Table S10:**
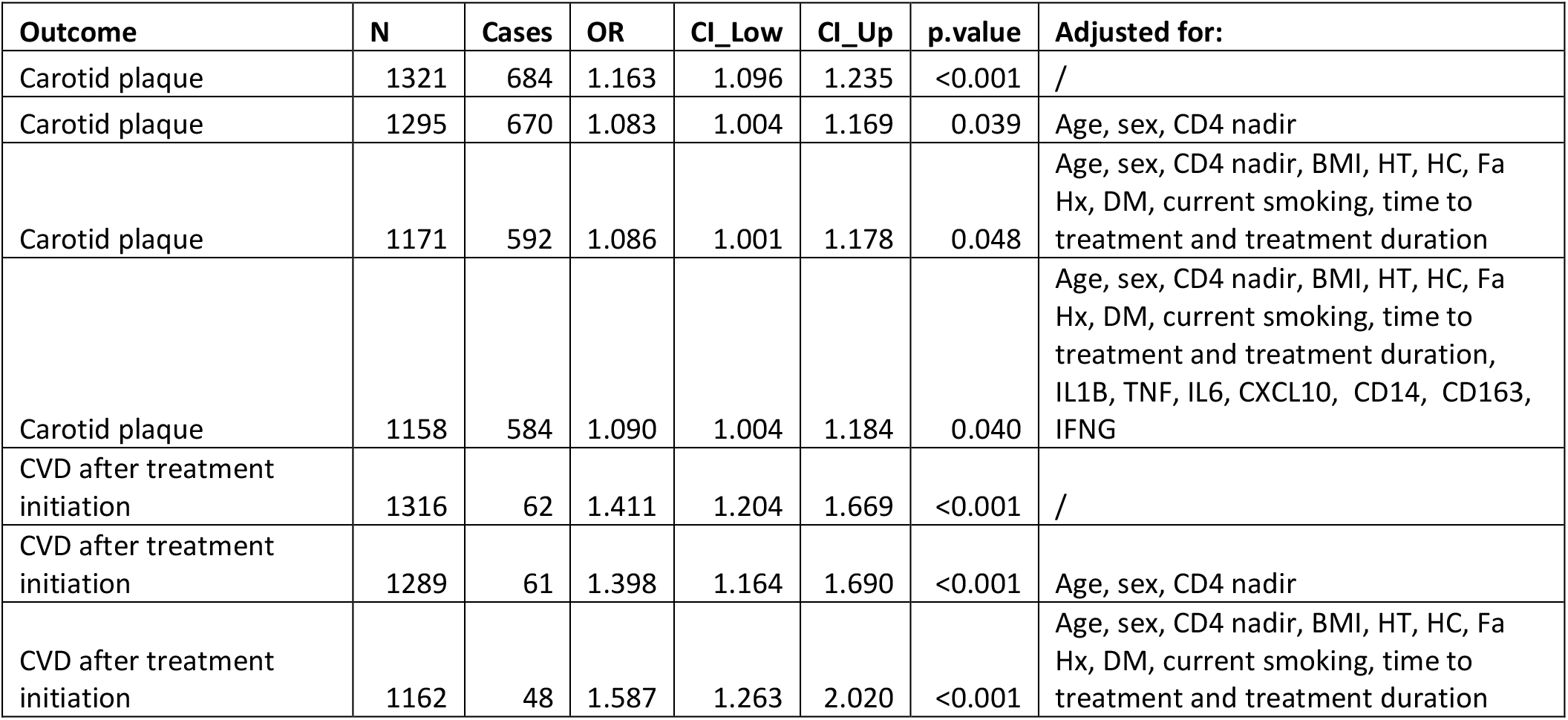
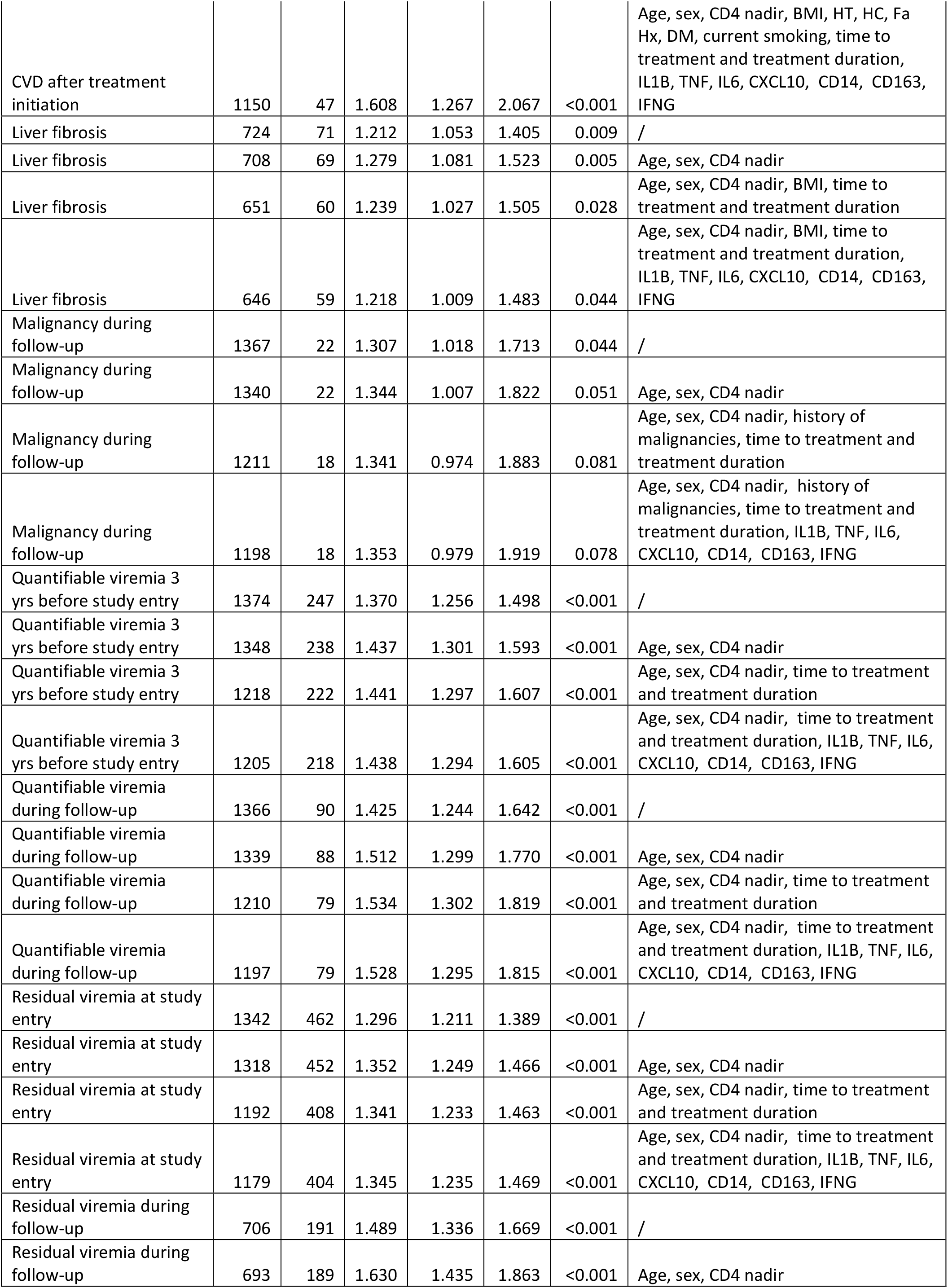
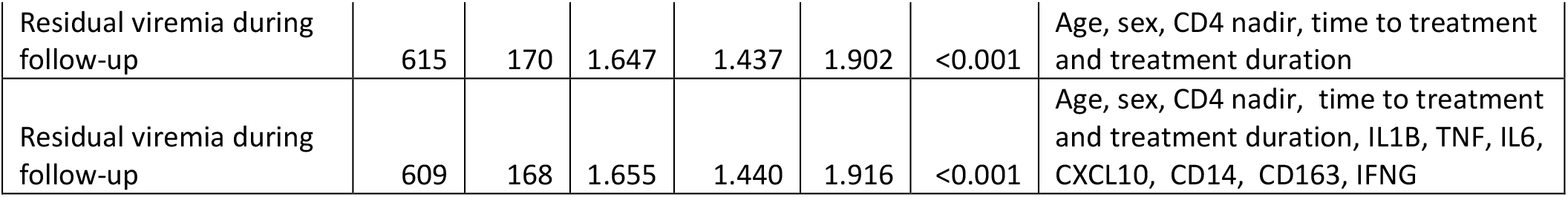
Univariable and mutlivariable models for different outcomes with defect HIV DNA as predictor. (N = sample size data in model, Cases = events of outcome, OR = odds ratio, CI_Low = lower limit 95%CI, CI_Up = upper limit 95%CI)

| **Outcome** | **N** | **Cases** | **OR** | **CI_Low** | **CI_Up** | **p.value** | **Adjusted for:** |
|---|---|---|---|---|---|---|---|
| Carotid plaque | 1321 | 684 | 1.163 | 1.096 | 1.235 | <0.001 | / |
| Carotid plaque | 1295 | 670 | 1.083 | 1.004 | 1.169 | 0.039 | Age, sex, CD4 nadir |
| Carotid plaque | 1171 | 592 | 1.086 | 1.001 | 1.178 | 0.048 | Age, sex, CD4 nadir, BMI, HT, HC, Fa Hx, DM, current smoking, time to treatment and treatment duration |
| Carotid plaque | 1158 | 584 | 1.090 | 1.004 | 1.184 | 0.040 | Age, sex, CD4 nadir, BMI, HT, HC, Fa Hx, DM, current smoking, time to treatment and treatment duration, IL1B, TNF, IL6, CXCL10, CD14, CD163, IFNG |
| CVD after treatment initiation | 1316 | 62 | 1.411 | 1.204 | 1.669 | <0.001 | / |
| CVD after treatment initiation | 1289 | 61 | 1.398 | 1.164 | 1.690 | <0.001 | Age, sex, CD4 nadir |
| CVD after treatment initiation | 1162 | 48 | 1.587 | 1.263 | 2.020 | <0.001 | Age, sex, CD4 nadir, BMI, HT, HC, Fa Hx, DM, current smoking, time to treatment and treatment duration |
| CVD after treatment initiation | 1150 | 47 | 1.608 | 1.267 | 2.067 | <0.001 | Age, sex, CD4 nadir, BMI, HT, HC, Fa Hx, DM, current smoking, time to treatment and treatment duration, IL1B, TNF, IL6, CXCL10, CD14, CD163, IFNG |
| Liver fibrosis | 724 | 71 | 1.212 | 1.053 | 1.405 | 0.009 | / |
| Liver fibrosis | 708 | 69 | 1.279 | 1.081 | 1.523 | 0.005 | Age, sex, CD4 nadir |
| Liver fibrosis | 651 | 60 | 1.239 | 1.027 | 1.505 | 0.028 | Age, sex, CD4 nadir, BMI, time to treatment and treatment duration |
| Liver fibrosis | 646 | 59 | 1.218 | 1.009 | 1.483 | 0.044 | Age, sex, CD4 nadir, BMI, time to treatment and treatment duration, IL1B, TNF, IL6, CXCL10, CD14, CD163, IFNG |
| Malignancy during follow-up | 1367 | 22 | 1.307 | 1.018 | 1.713 | 0.044 | / |
| Malignancy during follow-up | 1340 | 22 | 1.344 | 1.007 | 1.822 | 0.051 | Age, sex, CD4 nadir |
| Malignancy during follow-up | 1211 | 18 | 1.341 | 0.974 | 1.883 | 0.081 | Age, sex, CD4 nadir, history of malignancies, time to treatment and treatment duration |
| Malignancy during follow-up | 1198 | 18 | 1.353 | 0.979 | 1.919 | 0.078 | Age, sex, CD4 nadir, history of malignancies, time to treatment and treatment duration, IL1B, TNF, IL6, CXCL10, CD14, CD163, IFNG |
| Quantifiable viremia 3 yrs before study entry | 1374 | 247 | 1.370 | 1.256 | 1.498 | <0.001 | / |
| Quantifiable viremia 3 yrs before study entry | 1348 | 238 | 1.437 | 1.301 | 1.593 | <0.001 | Age, sex, CD4 nadir |
| Quantifiable viremia 3 yrs before study entry | 1218 | 222 | 1.441 | 1.297 | 1.607 | <0.001 | Age, sex, CD4 nadir, time to treatment and treatment duration |
| Quantifiable viremia 3 yrs before study entry | 1205 | 218 | 1.438 | 1.294 | 1.605 | <0.001 | Age, sex, CD4 nadir, time to treatment and treatment duration, IL1B, TNF, IL6, CXCL10, CD14, CD163, IFNG |
| Quantifiable viremia during follow-up | 1366 | 90 | 1.425 | 1.244 | 1.642 | <0.001 | / |
| Quantifiable viremia during follow-up | 1339 | 88 | 1.512 | 1.299 | 1.770 | <0.001 | Age, sex, CD4 nadir |
| Quantifiable viremia during follow-up | 1210 | 79 | 1.534 | 1.302 | 1.819 | <0.001 | Age, sex, CD4 nadir, time to treatment and treatment duration |
| Quantifiable viremia during follow-up | 1197 | 79 | 1.528 | 1.295 | 1.815 | <0.001 | Age, sex, CD4 nadir, time to treatment and treatment duration, IL1B, TNF, IL6, CXCL10, CD14, CD163, IFNG |
| Residual viremia at study entry | 1342 | 462 | 1.296 | 1.211 | 1.389 | <0.001 | / |
| Residual viremia at study entry | 1318 | 452 | 1.352 | 1.249 | 1.466 | <0.001 | Age, sex, CD4 nadir |
| Residual viremia at study entry | 1192 | 408 | 1.341 | 1.233 | 1.463 | <0.001 | Age, sex, CD4 nadir, time to treatment and treatment duration |
| Residual viremia at study entry | 1179 | 404 | 1.345 | 1.235 | 1.469 | <0.001 | Age, sex, CD4 nadir, time to treatment and treatment duration, IL1B, TNF, IL6, CXCL10, CD14, CD163, IFNG |
| Residual viremia during follow-up | 706 | 191 | 1.489 | 1.336 | 1.669 | <0.001 | / |
| Residual viremia during follow-up | 693 | 189 | 1.630 | 1.435 | 1.863 | <0.001 | Age, sex, CD4 nadir |
| Residual viremia during follow-up | 615 | 170 | 1.647 | 1.437 | 1.902 | <0.001 | Age, sex, CD4 nadir, time to treatment and treatment duration |
| Residual viremia during follow-up | 609 | 168 | 1.655 | 1.440 | 1.916 | <0.001 | Age, sex, CD4 nadir, time to treatment and treatment duration, IL1B, TNF, IL6, CXCL10, CD14, CD163, IFNG |

**Table S11:** Univariable and mutlivariable models for different outcomes with total HIV RNA as predictor. (N = sample size data in model, Cases = events of outcome, OR = odds ratio, CI_Low = lower limit 95%CI, CI_Up = upper limit 95%CI)

| Outcome | N | Cases | OR | CI_Low | CI_Up | p.value | Adjusted for: |
|---|---|---|---|---|---|---|---|
| Carotid plaque | 1361 | 692 | 1.074 | 1.032 | 1.118 | <0.001 | / |
| Carotid plaque | 1337 | 679 | 1.008 | 0.959 | 1.058 | 0.759 | Age, sex, CD4 nadir |
| Carotid plaque | 1209 | 598 | 1.012 | 0.960 | 1.067 | 0.658 | Age, sex, CD4 nadir, BMI, HT, HC, Fa Hx, DM, current smoking, time to treatment and treatment duration |
| Carotid plaque | 1197 | 591 | 1.015 | 0.962 | 1.070 | 0.597 | Age, sex, CD4 nadir, BMI, HT, HC, Fa Hx, DM, current smoking, time to treatment and treatment duration, IL1B, TNF, IL6, CXCL10, CD14, CD163, IFNG |
| CVD after treatment initiation | 1341 | 61 | 1.200 | 1.083 | 1.337 | 0.001 | / |
| CVD after treatment initiation | 1316 | 60 | 1.171 | 1.043 | 1.323 | 0.010 | Age, sex, CD4 nadir |
| CVD after treatment initiation | 1185 | 47 | 1.180 | 1.023 | 1.374 | 0.028 | Age, sex, CD4 nadir, BMI, HT, HC, Fa Hx, DM, current smoking, time to treatment and treatment duration |
| CVD after treatment initiation | 1173 | 46 | 1.170 | 1.012 | 1.368 | 0.040 | Age, sex, CD4 nadir, BMI, HT, HC, Fa Hx, DM, current smoking, time to treatment and treatment duration, IL1B, TNF, IL6, CXCL10, CD14, CD163, IFNG |
| Liver fibrosis | 724 | 70 | 1.109 | 1.010 | 1.221 | 0.033 | / |
| Liver fibrosis | 709 | 68 | 1.120 | 1.007 | 1.254 | 0.042 | Age, sex, CD4 nadir |
| Liver fibrosis | 652 | 60 | 1.091 | 0.971 | 1.235 | 0.154 | Age, sex, CD4 nadir, BMI, time to treatment and treatment duration |
| Liver fibrosis | 647 | 59 | 1.069 | 0.950 | 1.212 | 0.282 | Age, sex, CD4 nadir, BMI, time to treatment and treatment duration, IL1B, TNF, IL6, CXCL10, CD14, CD163, IFNG |
| Malignancy during follow-up | 1395 | 25 | 1.151 | 0.987 | 1.357 | 0.083 | / |
| Malignancy during follow-up | 1370 | 25 | 1.157 | 0.977 | 1.388 | 0.103 | Age, sex, CD4 nadir |
| Malignancy during follow-up | 1237 | 21 | 1.157 | 0.957 | 1.420 | 0.147 | Age, sex, CD4 nadir, history of malignancies, time to treatment and treatment duration |
| Malignancy during follow-up | 1225 | 21 | 1.206 | 0.981 | 1.513 | 0.090 | Age, sex, CD4 nadir, history of malignancies, time to treatment and treatment duration, IL1B, TNF, IL6, CXCL10, CD14, CD163, IFNG |
| Quantifiable viremia 3 yrs before study entry | 1399 | 267 | 1.133 | 1.076 | 1.196 | <0.001 | / |
| Quantifiable viremia 3 yrs before study entry | 1376 | 259 | 1.126 | 1.062 | 1.196 | <0.001 | Age, sex, CD4 nadir |
| Quantifiable viremia 3 yrs before study entry | 1241 | 240 | 1.149 | 1.078 | 1.226 | <0.001 | Age, sex, CD4 nadir, time to treatment and treatment duration |
| Quantifiable viremia 3 yrs before study entry | 1229 | 236 | 1.134 | 1.064 | 1.211 | <0.001 | Age, sex, CD4 nadir, time to treatment and treatment duration, IL1B, TNF, IL6, CXCL10, CD14, CD163, IFNG |
| Quantifiable viremia during follow-up | 1393 | 95 | 1.184 | 1.089 | 1.292 | <0.001 | / |
| Quantifiable viremia during follow-up | 1368 | 93 | 1.201 | 1.092 | 1.327 | <0.001 | Age, sex, CD4 nadir |
| Quantifiable viremia during follow-up | 1236 | 82 | 1.223 | 1.101 | 1.365 | <0.001 | Age, sex, CD4 nadir, time to treatment and treatment duration |
| Quantifiable viremia during follow-up | 1224 | 82 | 1.220 | 1.097 | 1.365 | <0.001 | Age, sex, CD4 nadir, time to treatment and treatment duration, IL1B, TNF, IL6, CXCL10, CD14, CD163, IFNG |
| Residual viremia at study entry | 1369 | 477 | 1.122 | 1.074 | 1.172 | <0.001 | / |
| Residual viremia at study entry | 1347 | 469 | 1.141 | 1.087 | 1.200 | <0.001 | Age, sex, CD4 nadir |
| Residual viremia at study entry | 1218 | 425 | 1.156 | 1.096 | 1.220 | <0.001 | Age, sex, CD4 nadir, time to treatment and treatment duration |
| Residual viremia at study entry | 1206 | 420 | 1.150 | 1.090 | 1.214 | <0.001 | Age, sex, CD4 nadir, time to treatment and treatment duration, IL1B, TNF, IL6, CXCL10, CD14, CD163, IFNG |
| Residual viremia during follow-up | 722 | 210 | 1.191 | 1.115 | 1.275 | <0.001 | / |
| Residual viremia during follow-up | 708 | 208 | 1.202 | 1.118 | 1.297 | <0.001 | Age, sex, CD4 nadir |
| Residual viremia during follow-up | 625 | 184 | 1.209 | 1.115 | 1.314 | <0.001 | Age, sex, CD4 nadir, time to treatment and treatment duration |
| Residual viremia during follow-up | 620 | 182 | 1.210 | 1.115 | 1.319 | <0.001 | Age, sex, CD4 nadir, time to treatment and treatment duration, IL1B, TNF, IL6, CXCL10, CD14, CD163, IFNG |

**Table S12:** Univariable and mutlivariable models for different outcomes with unspliced HIV RNA as predictor. (N = sample size data in model, Cases = events of outcome, OR = odds ratio, CI_Low = lower limit 95%CI, CI_Up = upper limit 95%CI)

| Outcome | N | Cases | OR | CI_Low | CI_Up | p.value | Adjusted for: |
|---|---|---|---|---|---|---|---|
| Carotid plaque | 1333 | 681 | 1.077 | 1.036 | 1.120 | <0.001 | / |
| Carotid plaque | 1310 | 669 | 1.008 | 0.960 | 1.058 | 0.751 | Age, sex, CD4 nadir |
| Carotid plaque | 1184 | 589 | 1.011 | 0.960 | 1.065 | 0.671 | Age, sex, CD4 nadir, BMI, HT, HC, Fa Hx, DM, current smoking, time to treatment and treatment duration |
| Carotid plaque | 1172 | 582 | 1.015 | 0.963 | 1.070 | 0.571 | Age, sex, CD4 nadir, BMI, HT, HC, Fa Hx, DM, current smoking, time to treatment and treatment duration, IL1B, TNF, IL6, CXCL10, CD14, CD163, IFNG |
| CVD after treatment initiation | 1314 | 59 | 1.205 | 1.097 | 1.327 | <0.001 | / |
| CVD after treatment initiation | 1290 | 58 | 1.185 | 1.064 | 1.325 | 0.002 | Age, sex, CD4 nadir |
| CVD after treatment initiation | 1161 | 45 | 1.219 | 1.070 | 1.398 | 0.004 | Age, sex, CD4 nadir, BMI, HT, HC, Fa Hx, DM, current smoking, time to treatment and treatment duration |
| CVD after treatment initiation | 1149 | 44 | 1.209 | 1.058 | 1.394 | 0.007 | Age, sex, CD4 nadir, BMI, HT, HC, Fa Hx, DM, current smoking, time to treatment and treatment duration, IL1B, TNF, IL6, CXCL10, CD14, CD163, IFNG |
| Liver fibrosis | 715 | 70 | 1.023 | 0.938 | 1.115 | 0.604 | / |
| Liver fibrosis | 700 | 68 | 1.013 | 0.920 | 1.116 | 0.787 | Age, sex, CD4 nadir |
| Liver fibrosis | 644 | 60 | 1.004 | 0.902 | 1.117 | 0.946 | Age, sex, CD4 nadir, BMI, time to treatment and treatment duration |
| Liver fibrosis | 639 | 59 | 0.980 | 0.877 | 1.095 | 0.727 | Age, sex, CD4 nadir, BMI, time to treatment and treatment duration, IL1B, TNF, IL6, CXCL10, CD14, CD163, IFNG |
| Malignancy during follow-up | 1367 | 25 | 1.136 | 0.986 | 1.311 | 0.079 | / |
| Malignancy during follow-up | 1343 | 25 | 1.123 | 0.961 | 1.317 | 0.147 | Age, sex, CD4 nadir |
| Malignancy during follow-up | 1212 | 21 | 1.092 | 0.919 | 1.303 | 0.321 | Age, sex, CD4 nadir, history of malignancies, time to treatment and treatment duration |
| Malignancy during follow-up | 1200 | 11 | 1.132 | 0.940 | 1.374 | 0.198 | Age, sex, CD4 nadir, history of malignancies, time to treatment and treatment duration, IL1B, TNF, IL6, CXCL10, CD14, CD163, IFNG |
| Quantifiable viremia 3 yrs before study entry | 1371 | 261 | 1.121 | 1.068 | 1.178 | <0.001 | / |
| Quantifiable viremia 3 yrs before study entry | 1349 | 253 | 1.120 | 1.060 | 1.184 | <0.001 | Age, sex, CD4 nadir |
| Quantifiable viremia 3 yrs before study entry | 1216 | 234 | 1.121 | 1.057 | 1.189 | <0.001 | Age, sex, CD4 nadir, time to treatment and treatment duration |
| Quantifiable viremia 3 yrs before study entry | 1204 | 230 | 1.110 | 1.047 | 1.178 | 0.001 | Age, sex, CD4 nadir, time to treatment and treatment duration, IL1B, TNF, IL6, CXCL10, CD14, CD163, IFNG |
| Quantifiable viremia during follow-up | 1365 | 95 | 1.094 | 1.015 | 1.179 | 0.019 | / |
| Quantifiable viremia during follow-up | 1341 | 93 | 1.093 | 1.006 | 1.190 | 0.037 | Age, sex, CD4 nadir |
| Quantifiable viremia during follow-up | 1211 | 82 | 1.108 | 1.012 | 1.214 | 0.027 | Age, sex, CD4 nadir, time to treatment and treatment duration |
| Quantifiable viremia during follow-up | 1199 | 82 | 1.117 | 1.019 | 1.226 | 0.020 | Age, sex, CD4 nadir, time to treatment and treatment duration, IL1B, TNF, IL6, CXCL10, CD14, CD163, IFNG |
| Residual viremia at study entry | 1341 | 468 | 1.080 | 1.037 | 1.125 | <0.001 | / |
| Residual viremia at study entry | 1320 | 460 | 1.086 | 1.038 | 1.137 | <0.001 | Age, sex, CD4 nadir |
| Residual viremia at study entry | 1193 | 416 | 1.098 | 1.045 | 1.155 | <0.001 | Age, sex, CD4 nadir, time to treatment and treatment duration |
| Residual viremia at study entry | 1181 | 411 | 1.094 | 1.040 | 1.151 | <0.001 | Age, sex, CD4 nadir, time to treatment and treatment duration, IL1B, TNF, IL6, CXCL10, CD14, CD163, IFNG |
| Residual viremia during follow-up | 703 | 202 | 1.101 | 1.038 | 1.169 | 0.002 | / |
| Residual viremia during follow-up | 690 | 200 | 1.098 | 1.028 | 1.173 | 0.006 | Age, sex, CD4 nadir |
| Residual viremia during follow-up | 609 | 177 | 1.112 | 1.035 | 1.197 | 0.004 | Age, sex, CD4 nadir, time to treatment and treatment duration |
| Residual viremia during follow-up | 604 | 175 | 1.122 | 1.041 | 1.210 | 0.003 | Age, sex, CD4 nadir, time to treatment and treatment duration, IL1B, TNF, IL6, CXCL10, CD14, CD163, IFNG |

**Table S13:**
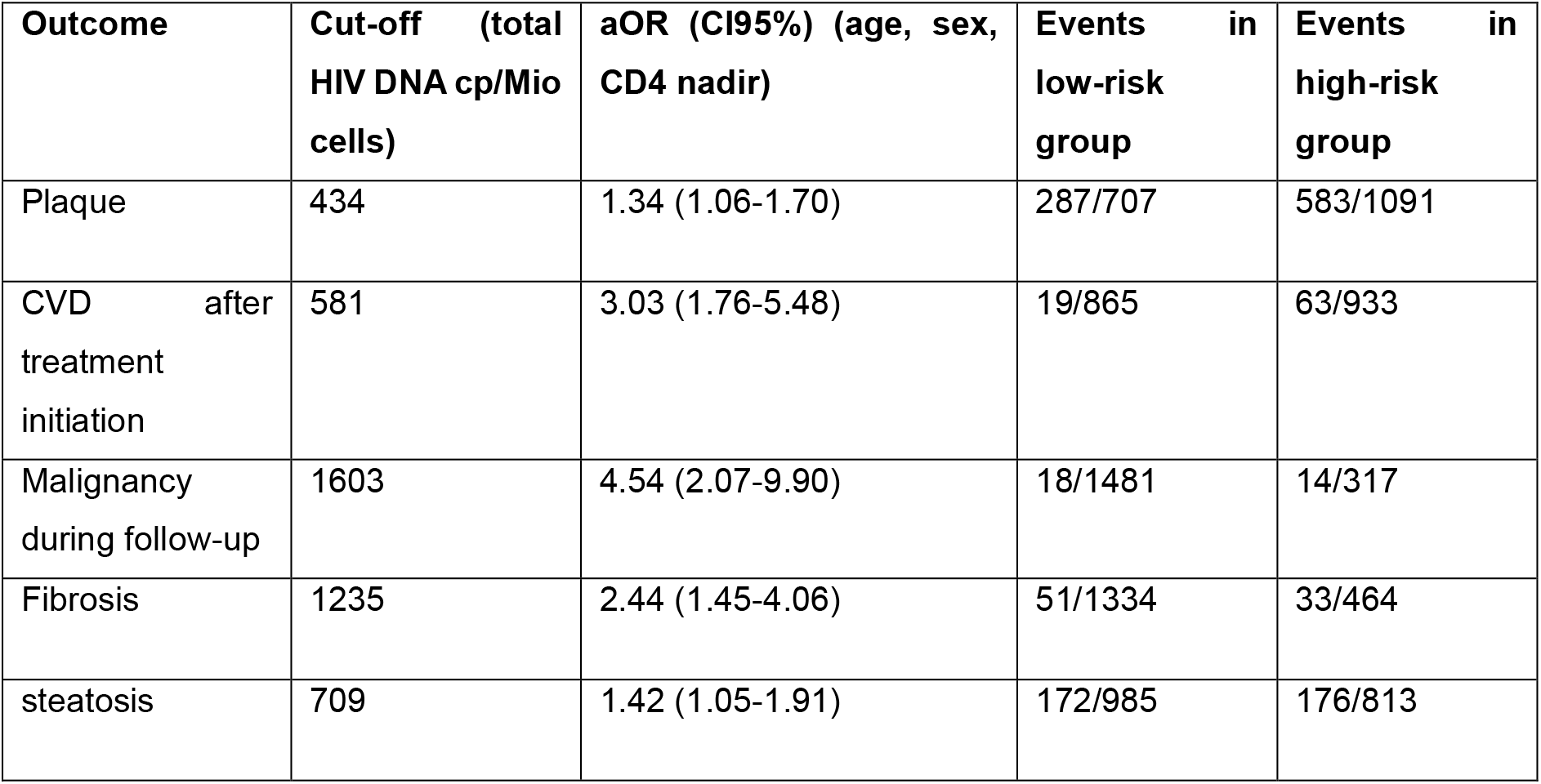
Cut-off level for each outcome as determined by the Youden index to maximize the balance between sensitivity and specificity. Odds ratio was adjusted for age, sex and CD4 nadir).

| **Outcome** | **Cut-off (total HIV DNA cp/Mio cells)** | **aOR (CI95%) (age, sex, CD4 nadir)** | **Events in low-risk group** | **Events in high-risk group** |
|---|---|---|---|---|
| Plaque | 434 | 1.34 (1.06-1.70) | 287/707 | 583/1091 |
| CVD after treatment initiation | 581 | 3.03 (1.76-5.48) | 19/865 | 63/933 |
| Malignancy during follow-up | 1603 | 4.54 (2.07-9.90) | 18/1481 | 14/317 |
| Fibrosis | 1235 | 2.44 (1.45-4.06) | 51/1334 | 33/464 |
| steatosis | 709 | 1.42 (1.05-1.91) | 172/985 | 176/813 |

## References

1. The Stages of HIV Infection. 2025 March 31, 2025; Available from: https://hivinfo.nih.gov/understanding-hiv/fact-sheets/stages-hiv-infection.

2. Jarrin, I., et al., Does rapid HIV disease progression prior to combination antiretroviral therapy hinder optimal CD4+ T-cell recovery once HIV-1 suppression is achieved? Aids, 2015. 29(17): p. 2323–33.

3. Jiang, C., et al., Distinct viral reservoirs in individuals with spontaneous control of HIV-1. Nature, 2020. 585(7824): p. 261–267.

4. Deeks, S.G., S.R. Lewin, and D.V. Havlir, The end of AIDS: HIV infection as a chronic disease. Lancet, 2013. 382(9903): p. 1525–33.

5. Lundgren, J.D., et al., Initiation of Antiretroviral Therapy in Early Asymptomatic HIV Infection. N Engl J Med, 2015. 373(9): p. 795–807.

6. Trickey, A., et al., Life expectancy after 2015 of adults with HIV on long-term antiretroviral therapy in Europe and North America: a collaborative analysis of cohort studies. Lancet HIV, 2023. 10(5): p. e295–e307.

7. Fomundam, H.N., et al., Prevalence and predictors of late presentation for HIV care in South Africa. S Afr Med J, 2017. 107(12): p. 1058–1064.

8. Parczewski, M., et al., Control of HIV across the WHO European region: progress and remaining challenges. Lancet Reg Health Eur, 2025. 52: p. 101243.

9. Schouten, J., et al., Cross-sectional comparison of the prevalence of age-associated comorbidities and their risk factors between HIV-infected and uninfected individuals: the AGEhIV cohort study. Clin Infect Dis, 2014. 59(12): p. 1787–97.

10. Battegay, M., et al., Immunological recovery and antiretroviral therapy in HIV-1 infection. Lancet Infect Dis, 2006. 6(5): p. 280–7.

11. Phan, T., et al., Understanding early HIV-1 rebound dynamics following antiretroviral therapy interruption: The importance of effector cell expansion. PLoS Pathog, 2024. 20(7): p. e1012236.

12. Hiener, B., et al., Identification of Genetically Intact HIV-1 Proviruses in Specific CD4(+) T Cells from Effectively Treated Participants. Cell Rep, 2017. 21(3): p. 813–822.

13. Siliciano, J.D., et al., Long-term follow-up studies confirm the stability of the latent reservoir for HIV-1 in resting CD4+ T cells. Nat Med, 2003. 9(6): p. 727–8.

14. Chun, T.W., et al., Presence of an inducible HIV-1 latent reservoir during highly active antiretroviral therapy. Proc Natl Acad Sci U S A, 1997. 94(24): p. 13193–7.

15. Chomont, N., et al., HIV reservoir size and persistence are driven by T cell survival and homeostatic proliferation. Nat Med, 2009. 15(8): p. 893–900.

16. Bruner, K.M., et al., Defective proviruses rapidly accumulate during acute HIV-1 infection. Nat Med, 2016. 22(9): p. 1043–9.

17. Linden, N. and R.B. Jones, Potential multi-modal effects of provirus integration on HIV-1 persistence: lessons from other viruses. Trends Immunol, 2022. 43(8): p. 617–629.

18. McLaughlin, M.M., et al., Association of Viral Persistence and Atherosclerosis in Adults With Treated HIV Infection. JAMA Netw Open, 2020. 3(10): p. e2018099.

19. Bachmann, N., et al., Determinants of HIV-1 reservoir size and long-term dynamics during suppressive ART. Nat Commun, 2019. 10(1): p. 3193.

20. González-Navarro, I., et al., Assessing advances in three decades of clinical antiretroviral therapy on the HIV-1 reservoir. J Clin Invest, 2024. 135(2).

21. Gasca-Capote, C., et al., The HIV-1 reservoir landscape in persistent elite controllers and transient elite controllers. J Clin Invest, 2024. 134(8).

22. Buzon, M.J., et al., Long-term antiretroviral treatment initiated at primary HIV-1 infection affects the size, composition, and decay kinetics of the reservoir of HIV-1-infected CD4 T cells. J Virol, 2014. 88(17): p. 10056–65.

23. Zhang, L.X., et al., Dynamics of HIV reservoir decay and naïve CD4 T-cell recovery between immune non-responders and complete responders on long-term antiretroviral treatment. Clin Immunol, 2021. 229: p. 108773.

24. Vos, W., et al., The 2000HIV study: Design, multi-omics methods and participant characteristics. Front Immunol, 2022. 13: p. 982746.

25. Otten, T., et al., Impact of COVID-19, lockdowns and vaccination on immune responses in a HIV cohort in the Netherlands. Front Immunol, 2024. 15: p. 1459593.

26. Zhang, Y., et al., Opposite effects of chronic HIV infection and antiretroviral medication on organismal and organ-specific biological aging. Nat Commun, 2026. 17(1).

27. Höft, M.A., W.A. Burgers, and C. Riou, The immune response to SARS-CoV-2 in people with HIV. Cell Mol Immunol, 2024. 21(2): p. 184–196.

28. Menzel, A., et al., Common and Novel Markers for Measuring Inflammation and Oxidative Stress Ex Vivo in Research and Clinical Practice-Which to Use Regarding Disease Outcomes? Antioxidants (Basel), 2021. 10(3).

29. Imamichi, H., et al., Defective HIV-1 proviruses produce novel protein-coding RNA species in HIV-infected patients on combination antiretroviral therapy. Proc Natl Acad Sci U S A, 2016. 113(31): p. 8783–8.

30. Imamichi, H., et al., Defective HIV-1 proviruses produce viral proteins. Proc Natl Acad Sci U S A, 2020. 117(7): p. 3704–3710.

31. Martin, H.A., et al., New Assay Reveals Vast Excess of Defective over Intact HIV-1 Transcripts in Antiretroviral Therapy-Suppressed Individuals. J Virol, 2022. 96(24): p. e0160522.

32. Zicari, S., et al., Immune Activation, Inflammation, and Non-AIDS Co-Morbidities in HIV-Infected Patients under Long-Term ART. Viruses, 2019. 11(3).

33. Lange, C.G., et al., CD4+ T-lymphocyte nadir and the effect of highly active antiretroviral therapy on phenotypic and functional immune restoration in HIV-1 infection. Clin Immunol, 2002. 102(2): p. 154–61.

34. White, J.A., et al., Complex decay dynamics of HIV virions, intact and defective proviruses, and 2LTR circles following initiation of antiretroviral therapy. Proc Natl Acad Sci U S A, 2022. 119(6).

35. Carty, M., C. Guy, and A.G. Bowie, Detection of Viral Infections by Innate Immunity. Biochem Pharmacol, 2021. 183: p. 114316.

36. Heil, F., et al., Species-specific recognition of single-stranded RNA via toll-like receptor 7 and 8. Science, 2004. 303(5663): p. 1526–9.

37. Rezagholizadeh, N., et al., TLR7 Mediates HIV-1 Tat-Induced Cellular Senescence in Human Astrocytes. Aging Cell, 2025. 24(7): p. e70086.

